# FLT3 is genetically essential for ITD-mutated leukemic stem cells but dispensable for human hematopoietic stem cells

**DOI:** 10.1101/2024.06.30.601363

**Authors:** Joana L Araujo, Elvin Wagenblast, Veronique Voisin, Jessica McLeod, Olga I. Gan, Suraj Bansal, Liqing Jin, Amanda Mitchell, Blaise Gratton, Sarah Cutting, Andrea Arruda, Monica Doedens, Jose-Mario Capo-Chichi, Sagi Abelson, Mark D Minden, Jean C. Y. Wang, Manuel A. Sobrinho-Simões, Perpétua Pinto-do-Ó, Eric Lechman, John E. Dick

## Abstract

Leukemic stem cells (LSCs) fuel acute myeloid leukemia (AML) growth and relapse, but therapies tailored towards eradicating LSCs without harming healthy hematopoietic stem cells (HSCs) are lacking. FLT3 is considered an important therapeutic target due to frequent mutation in AML and association with relapse. However, there has been limited clinical success with FLT3 targeting, suggesting either that FLT3 is not a vulnerability in LSC, or that more potent inhibition is required, a scenario where HSC toxicity could become limiting. We tested these possibilities by ablating FLT3 using CRISPR/Cas9-mediated FLT3 knock-out (FLT3-KO) in human LSCs and HSCs followed by functional xenograft assays. FLT3-KO in LSCs from FLT3-ITD mutated, but not FLT3-WT AMLs, resulted in short-term leukemic grafts of FLT-3-KO edited cells that disappeared by 12 weeks. By contrast, FLT3-KO in HSCs from fetal liver, cord blood and adult bone marrow did not impair multilineage hematopoiesis in primary and secondary xenografts. Our study establishes FLT3 as an ideal therapeutic target where ITD+ LSC are eradicated upon FLT3 deletion, while HSCs are spared. These findings support the development of more potent FLT3-targeting drugs or gene-editing approaches for LSC eradication to improve clinical outcomes.

**KEY POINTS:** The FLT3 gene is essential for ITD-mutated leukemic stem cells (LSCs) to establish and propagate leukemia.

Normal human hematopoietic stem cells (HSCs) do not require FLT3 to engraft and sustain hematopoiesis.

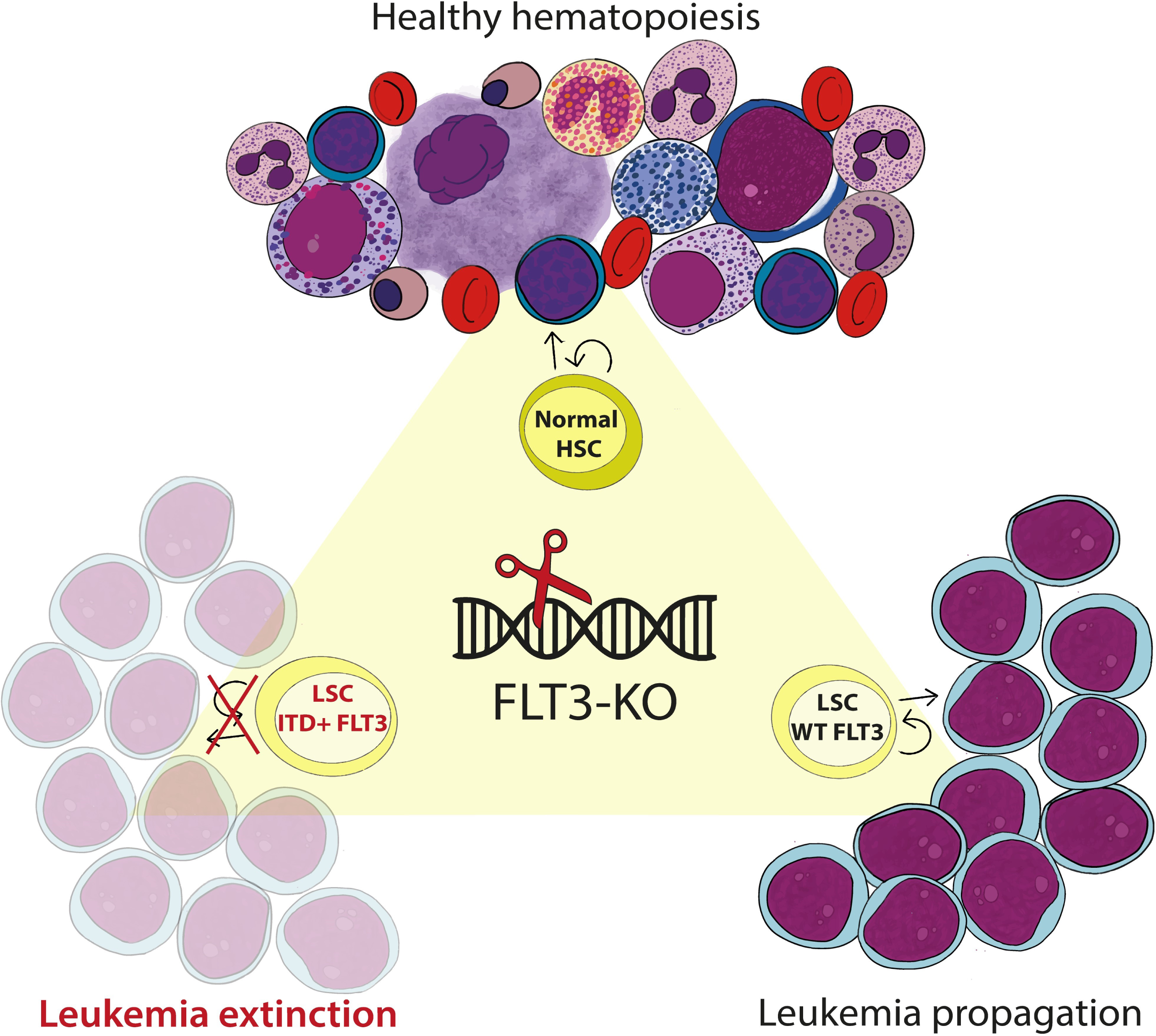

## INTRODUCTION

Acute myeloid leukemia (AML) is a caricature of normal blood development^1^ with leukemic stem cells (LSCs) residing at the top of the malignant hierarchy, resisting chemotherapy and serving as a reservoir for relapse^2–5^. As most AML patients die of relapse^6^, better targeting of LSCs is required, but LSCs and hematopoietic stem cells (HSCs) share numerous biological properties, raising the question of how to intensively target LSCs without harming healthy HSCs.

FMS-like Tyrosine Kinase 3 (FLT3) is an appealing therapeutic target as it is a powerful oncogene and the most frequently mutated gene in AML^7–9^. The internal tandem duplication (ITD) of FLT3 is an activating mutation linked to increased chemoresistance, relapse and mortality, even following allogeneic stem cell transplantation^10–12^. Detection of FLT3-ITD mutation in morphological remission is reported to be a stronger predictor of relapse and death than methods currently used to detect measurable residual disease^13^. The human FLT3 gene was cloned in 1993^14^, activating mutations identified in 1996^15^ and clinical studies with FLT3 inhibitors started in 2000^16^. However, many years passed before FLT3 inhibitors produced results significant enough to justify their widespread clinical use^17^. Despite consistent progress^18^, the use of FLT3 inhibitors in frontline or relapsed FLT3-mutant AML has only met with modest success^17,19,20^. Key issues over the years have been the lack of selectivity and potency of inhibitors, the limited duration of inhibition, occurrence of resistance-conferring FLT3 mutations and clonal adaptation that drives resistance to FLT3 targeting. While deeper inhibition or even deletion of FLT3 might overcome some of these issues, it is still unknown if LSCs are dependent on FLT3. Moreover, clinical efficacy has two requirements: that LSC are eradicated to prevent recurrence and that normal HSCs are able to tolerate full FLT3 inhibition. There have been no comprehensive mechanistic and functional studies to determine whether FLT3 is essential for human HSCs or LSCs. The few available human studies do suggest that FLT3 is expressed in HSCs^21–24^, however its functional importance has not been reported. In the mouse, FLT3 does mark important functional differences among hematopoietic progenitors, however it is well known that HSCs do not express FLT3^25–30^. This divergence in FLT3 expression on murine and human HSCs renders mouse models unsuitable for exploring the role of FLT3 in human hematopoiesis.

Here, we performed CRISPR/Cas9-mediated knock-out (KO) on human hematopoietic populations to evaluate the effects of FLT3 deletion in human LSCs and healthy HSCs across ontogeny. Using functional and transcriptomic studies, we demonstrate that FLT3 is genetically essential for LSCs to drive leukemogenesis in FLT3-ITD AML but is dispensable for both FLT3-WT AML and healthy HSC-driven hematopoiesis.

## METHODS

Detailed experimental methods are described in Supplemental Methods.

### Human primary samples collection and cell sorting

Fetal liver (FL) samples were collected from elective pregnancy terminations at 16-19 weeks of gestation at Mount Sinai Hospital, processed within 3 hours (h) and CD34+ cells were isolated (Miltenyi Biotec) and viably frozen at -150°C. Umbilical cord blood (CB) samples were collected at Trillium, William Osler and Credit Valley Hospitals, processed within 24-48h after birth, lineage (Lin)-positive cells depletion was performed (StemCell Technologies) and Lin-cells were viably frozen. Adult bone marrow (BM) samples were collected during elective hip replacement at Centro Hospitalar Universitário de São João, and mononuclear cells were viably frozen. Primary AML samples were obtained from patients’ peripheral blood at Princess Margaret Cancer Centre, and mononuclear cells were viably frozen. All human samples were collected after patient informed written consent and in accordance with guidelines approved by the ethics board from each mentioned health institution and University Health Network (UHN) Research Ethics Board. After thawing, HSCs defined as CD34+CD38-CD45RA-CD90+CD49f+ were sorted from FL and CB, and hematopoietic stem and progenitor cells (HSPCs) defined as HSCs and MPPs (CD34+CD38-CD45RA-CD90-CD49f-) were sorted from BM. AML samples (Table 1) were sorted according to previously functionally validated LSC-enriched populations (see supplemental methods): CD34+ leukemic cells were sorted from all AML samples except for WT3, for which CD45+CD3-population was sorted.

**Table 1.** Characterization of AML samples. Targeted next-generation sequencing to characterize the mutational composition of the samples was obtained using Single-molecule molecular inversion probes (smMIPs)-based NGS tools described in Medeiros 2022 (samples labeled with “a”); or was already available in the patients included in the Advanced Genomics in Leukemia (AGILE) prospective trial from Princess Margaret Cancer Centre (Kamel-Reid, 2015) (samples labeled with “b”). FLT3-TKD mutation was not detected in any sample. Samples ITD 5, ITD 6 and ITD 7 were obtained from patients that did not respond to chemotherapy combined with midostaurin (Midostaurin-R).

| Sample ID | FLT3 Status | Other mutations | Karyotype | ELN2022 Risk | Disease stage | Notes |
| --- | --- | --- | --- | --- | --- | --- |
| **ITD 1<sup>a</sup>** | ITD | WT1, ASXL1, U2AF1, KRAS, BCOR. | Normal | Adverse | Refractory |  |
| **ITD 2<sup>a</sup>** | ITD | NPM1, DNMT3A. | Normal | Intermediate | Diagnosis |  |
| **ITD 3<sup>a</sup>** | ITD | PHF6, WT1. | Complex | Adverse | Relapse |  |
| **ITD 4<sup>a</sup>** | ITD | NPM1, DNMT3A. | Normal | Intermediate | Diagnosis |  |
| **ITD 5<sup>b</sup>** | ITD | NPM1, WT1. | Normal | Intermediate | Diagnosis | Midostaurin-R |
| **ITD 6<sup>b</sup>** | ITD | NPM1, DNMT3A. | Failure | Intermediate | Relapse | Midostaurin-R |
| **ITD 7<sup>b</sup>** | ITD | RUNX1, U2AF1. | Del20q | Adverse | Diagnosis | Midostaurin-R |
| **WT 1<sup>a</sup>** | WT | NRAS, WT1, CEBPA. | Normal | Intermediate | Relapse |  |
| **WT 2<sup>a</sup>** | WT | NRAS, TP53, SF3B1. | Complex | Adverse | Relapse |  |
| **WT 3<sup>a</sup>** | WT | TET2, CBL. | Complex | Adverse | Diagnosis |  |

a - NGS by smMIP panel; b - NGS by AGILE panel

### FLT3-surface expression

A 2-step staining was optimized to characterize FLT3 surface expression and to perform cell-sorting according to FLT3 expression. Cells were primarily stained with FLT3 CD135 biotin-conjugated mouse anti-human CD135, clone 4G8 (BD, 1:50), washed twice and subsequently stained with streptavidin PE (BD, 1:250), all steps at 4°C.

### CRISPR/Cas9-mediated FLT3-KO on *in vivo* assays

To edit FLT3, a pair of gRNAs deleting 88 base pairs at the exon 20 of FLT3 gene was selected from 10 predicted by CRoatan algorithm (http://croatan.hannonlab.org/). Control gene (the olfactory receptor OR2W5) was edited by a pair of gRNAs previously validated^31^. FL, CB HSCs and BM HSPCs were cultured for 48h and AML cells for 24h and then CRISPR/Cas9 RNP electroporation was performed using the 4D-Nucleofector (Lonza), chemically synthesized gRNAs (IDT) and recombinant Cas9 nuclease (IDT)^32,33^. After recovering overnight, cells were intrafemorally injected into sublethally irradiated NSG female mice, except for experiments where alternative recipients were used (see supplemental methods). Short-term (2 weeks), long term (12 to 20 weeks), primary and secondary xenotransplantation studies were performed. All mouse experiments were approved by the Animal Care Committee of the UHN. After euthanizing the mice, injected femur and non-injected bones (contralateral femur and tibias) were flushed separately so bone marrow was independently collected and analyzed by flow cytometry to characterize human hematopoietic engraftment, and genotyping and cyto-morphological analysis were performed. Spleens were collected, crushed and similarly analyzed. In some instances, spleens were fixated and embedded in paraffin for subsequent histological analysis. Percentage of gene-KO was determined in DNA from cells collected before transplantation and from engrafted cells, by Sanger sequencing and indel analysis using the online tool ICE Synthego^34^.

### Competition assay and RNA sequencing

CD34+ cells were sorted from sample ITD3, and CD34+38-cells were sorted from CB, FLT3-KO and OR2W5-KO were performed in each sample individually, and both cell-populations were co-transplanted in a 1:1 proportion using 10 000-20 000 cells from each sample per mouse. Human engraftment was analysed to determine the presence and proportion of multilineage hematopoiesis *versus* leukemia, using flow cytometry, cyto-morphology, histology and genotyping analysis in primary and secondary recipients.

### RNA sequencing

Three samples from each condition – ITD+ AML (ITD1-3), AML without ITD mutation (WT1-3) and CB (3 samples) – were sorted: CD34+ population from all AML samples, except WT3 in which CD45+CD3-population was sorted, and CD34+38-population from CB samples. After a 24-48h culture, RNA was isolated using the RNeasy Micro kit (Qiagen) and deep sequencing was performed on the NovaSeq 6000 system. Differential gene-expression and pathway enrichment analysis were performed as described previously^35^.

## RESULTS

### 1. FLT3-KO impairs LSC function in FLT3-ITD but not FLT3-WT AML

To assess the importance of FLT3 expression for LSC function, we induced CRISPR/Cas9-mediated FLT3-KO in LSC-enriched populations from primary AML samples with FLT3-ITD mutation (ITD 1 to 7) and without FLT3-ITD mutation (WT 1 to 3) and tested their capacity to generate leukemia in NSG mice (**Fig. 1A, Table 1**). Most leukemic blasts and CD34+ leukemic cells in both ITD and WT AMLs expressed surface FLT3 at baseline (**Supp.** Fig. 1A). FLT3-KO was designed to target exon 20, affecting both ITD-mutated and WT alleles and resulting in an 88 base-pair frame-shift deletion near the catalytically active site (**Supp.** Fig. 1B and C). FLT3-KO significantly reduced the sizes of leukemic grafts and spleens in 6 out of 7 ITD+ leukemias compared to control-gene KO (targeting olfactory receptor OR2W5) (**Fig 1B, Supp.** Fig. 1D and E). By contrast, little to no effect on leukemic engraftment was seen in FLT3 WT leukemias (**Fig 1B, Supp.** Fig. 1D and E). FLT3 expression was drastically reduced in 12-week FLT3-KO grafts in both ITD (ITD7) and WT (WT2) AMLs compared with the control group (**Supp Fig 1F**), confirming that FLT3-KO disrupted protein expression.

**Figure 1.**
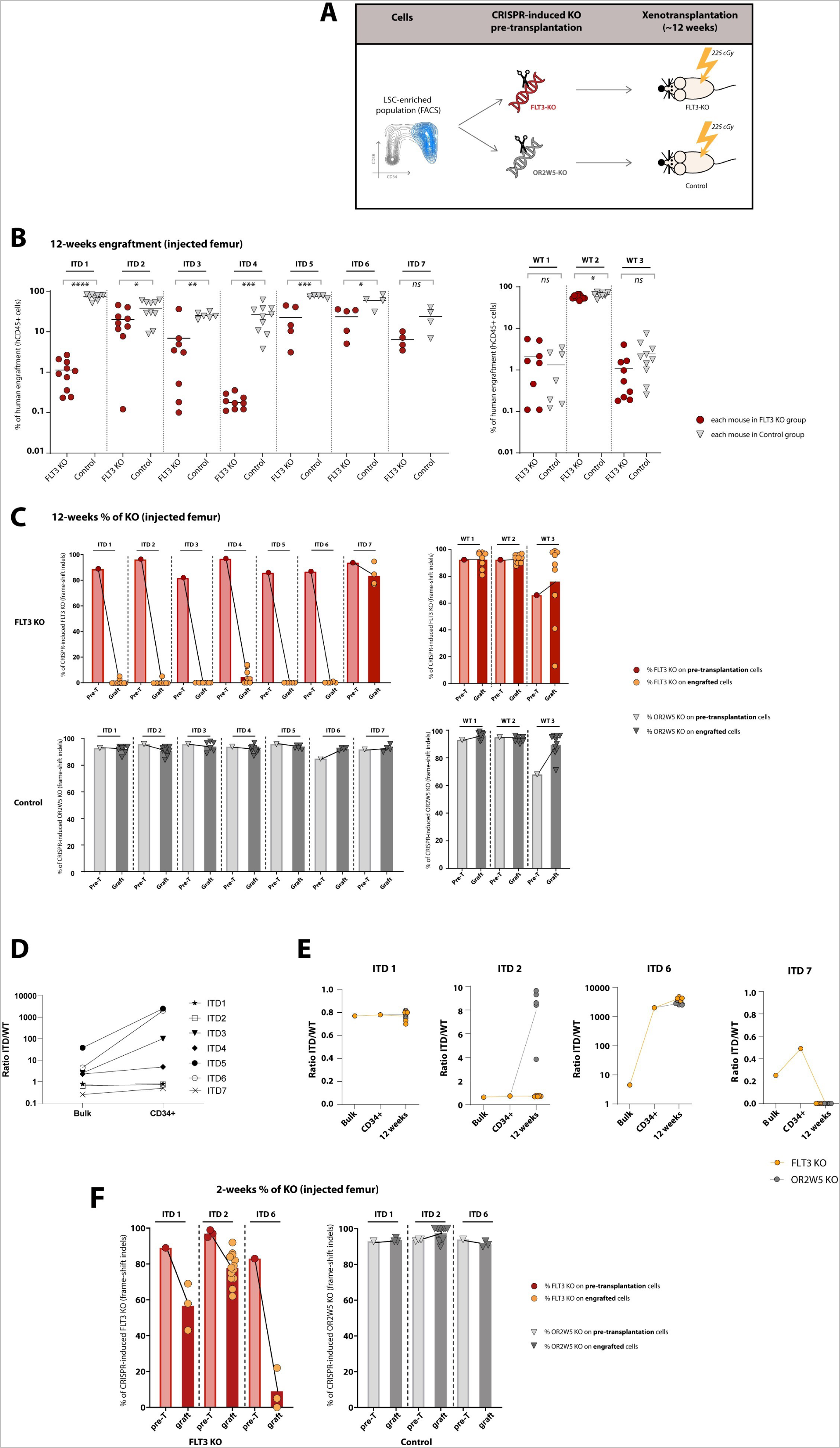
FLT3-KO prevents engraftment of FLT3-ITD LSCs but not of FLT3-WT LSCs. **(A)** Overview of the *in vivo* assay of FLT3-KO in AML. **(B)** Human engraftment of leukemic samples at 12 weeks, in sub-lethally irradiated NSG female mice injected with FLT3-KO or control gene (OR2W5)-KO cells; the experiment with sample ITD 7 was humanely terminated at 8 weeks; 4-10 mice per condition; results refer to the percentages of engraftment in the femurs that were originally injected with the cells (injected femur). **(C)** KO percentage in the cells pre-transplantation (pre-T) and in the engrafted (graft) cells from the FLT3-KO group (top) and control group (bottom), corresponding to the experiment in (B), determined by Sanger sequencing and indel analysis. **(D)** FLT3-ITD allele ratio (ITD/WT) determined by PCR and capillary electrophoresis in AML samples with FLT3-ITD mutation (ITD 1 to 7), in the bulk sample and in CD34+ sorted population. **(E)** FLT3-ITD allele ratio (ITD/WT) in AML samples ITD1, 2, 6 and 7, in the bulk sample, CD34+ sorted population and 12-week engrafted samples. **(F)** Short-term (2 weeks) transplants of ITD-mutated AMLs (ITD1, 2 and 6) in sublethally irradiated NSG female mice, KO percentage in the cells pre-transplantation (pre-T) and in the engrafted (graft) cells from FLT3 KO (left) and control groups (right), determined by Sanger sequencing and indel analysis; experiment with ITD2 was performed 3 times, whereas experiments with ITD1 and ITD were performed once; results refer to the injected femur; in the engrafted samples (graft) each symbol represents 1 mouse; engraftment levels are depicted in Supp. Fig 1H. Positive engraftment was considered if ≥0.1% human cells; lineage characterization was performed only on grafts with ≥1% of human cells. Unpaired Student’s t test: *P < 0.05; **P < 0.01; ***P < 0.001; ****P < 0.0001; mean ± standard deviation values are reported in the graphs.

To assess the degree of FLT3-KO editing in leukemic grafts, we determined the KO percentage in cells collected before transplantation and at 12 weeks post-transplant. In pre-transplantation cells, the percentage of cells with KO was greater than 80% in control and FLT3-KO groups (**Fig 1C**), establishing that all mice were transplanted with cells containing efficiently edited DNA. In 12-week leukemic grafts from samples ITD 1 to 6, the percentage of cells with FLT3-KO dropped close to zero, demonstrating that the small grafts were made up of cells that had escaped FLT3 gene editing (**Fig 1C**). FLT3-KO using a different pair of gRNAs in samples ITD 1 and 2 generated similar results (**Supp Fig 1G**), independently confirming our findings. By contrast, ITD7 and all WT samples maintained FLT3-KO in greater than 80% of cells within 12-week grafts, similar to controls (**Fig 1C**). Of all the ITD-mutated AML samples tested, ITD7 had the lowest ITD/WT ratio at baseline (**Fig. 1D**) and the ratio decreased to zero following transplantation, with no detectable ITD-mutation in engrafted cells, contrasting with samples ITD 1, 2 and 6, where the ITD/WT ratio was maintained or increased after transplantation (**Fig. 1E**). These findings reveal that sample ITD7 must have contained genetic subclones and that engraftment was driven by a FLT3 WT subclone rather than the ITD-mutated clone, thereby explaining why this sample behaved like a WT sample upon FLT3-KO.

To test whether FLT3-KO prevents early engraftment or leads to the elimination of ITD+ leukemic cells over-time, we induced FLT3-KO in samples ITD 1, 2 and 6 and performed short-term xenotransplants. Grafts were detectable at 2 weeks (**Supp.** Fig. 1H) and human DNA with FLT3-KO was detected at variable proportions (average 9 to 80%) (**Fig 1F**), but there was almost complete absence of FLT3-KO DNA at 12 weeks in the same samples (**Fig 1C**). These observations suggest that FLT3 is not needed for short-term engraftment of ITD+ leukemic progenitors but is required for the function of the rare self-renewing LSCs that sustain the ITD+ AML.

### 2. FLT3 is not required for repopulation and self-renewal in healthy human HSCs

To evaluate the requirement for FLT3 in normal human hematopoiesis, we first characterized FLT3 expression in human FL, CB and adult BM by cell surface and transcriptomic analysis. FLT3 was expressed in most HSCs and immature progenitors and became downregulated upon differentiation (**Fig 2A** and **Supp.** Fig 2A). HSCs expressing high or low levels of FLT3 produced equivalent grafts at 20 weeks in NSG mice (**Supp.** Fig 2B).

**Figure 2.**
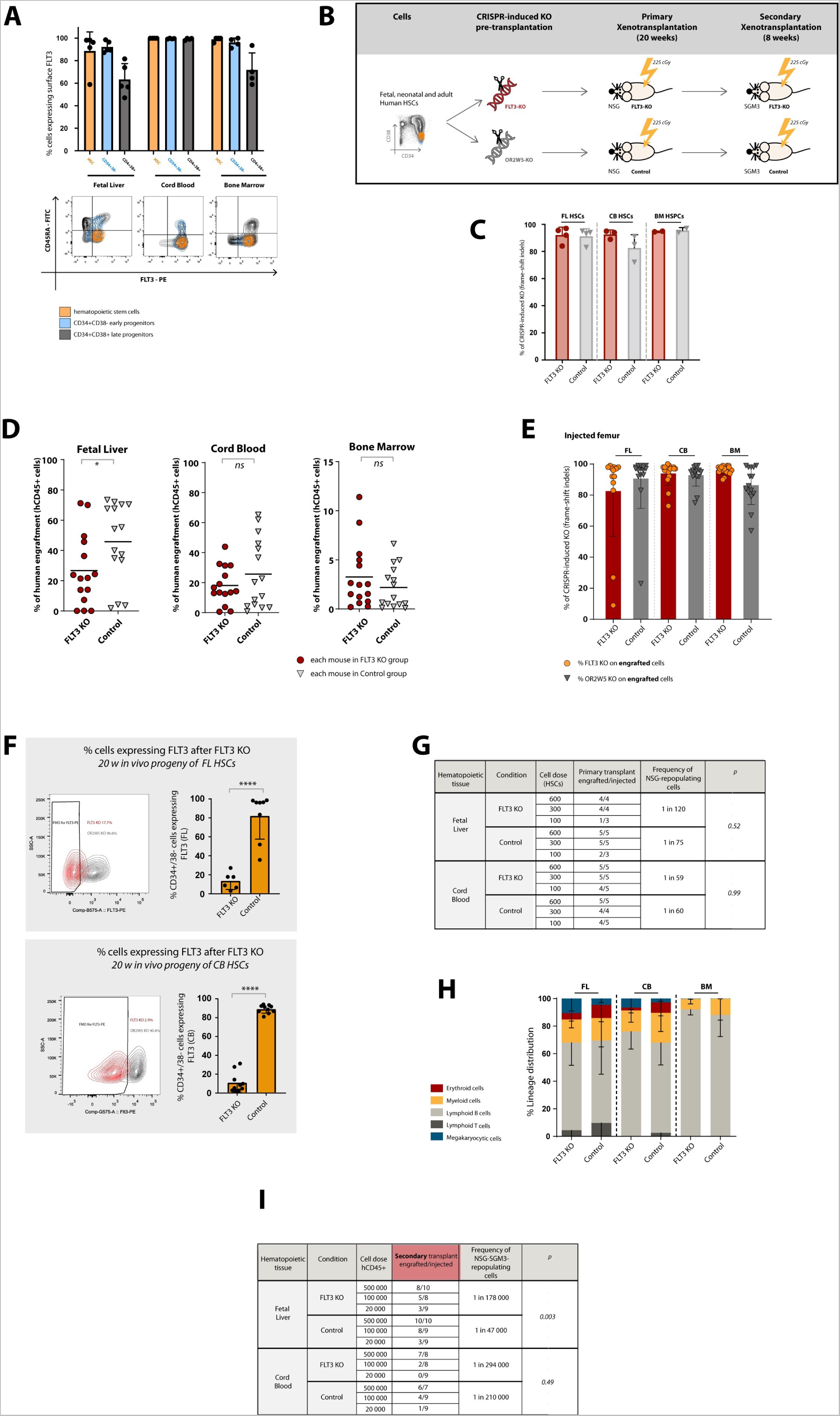
FLT3-KO in normal human HSCs does not prevent engraftment and self-renewal. **(A)** FLT3 cell-surface expression on HSCs (CD34+CD38-CD45RA-CD90+CD49f+), primitive progenitors (CD34+CD38-) and maturing progenitors (CD34+CD38+) from human FL, CB and adult BM, n=4 for each tissue. **(B)** Overview of the *in vivo* experiments of FLT3-KO in normal FL, CB and BM. **(C)** KO percentages of CRISPR/Cas9-mediated FLT3-KO and control gene-KO in human HSCs from FL (4 samples) and CB (3 samples) and in HSPCs (HSCs and MPPs defined as CD34+38-45RA-90-49f-) from BM (2 samples), determined by Sanger sequencing and indel analysis. **(D)** Levels of human engraftment in mice injected with FLT3-KO and control gene-KO HSCs from FL and CB and HSPCs from BM, at 20 weeks (FL, CB) and 12 weeks (BM); recipients were sublethally irradiated female NSG mice; percentages of engraftment in injected femurs; 3 human samples of each tissue were used in 3 independent experiments (15 mice per condition). **(E)** FLT3 and control gene-KO percentages in human cells engrafted in each mouse from (D); KO percentages were determined independently in the injected femurs; 2 mice transplanted with FLT3-KO FL HSCs were not engrafted by human cells and human DNA was not detectable, so genotyping was not performed on those. **(F)** FLT3 cell-surface expression (by flow cytometry) in CD34+CD38-human cells engrafted in mice from (D) injected with FL and CB cells. **(G)** NSG-repopulating cell frequencies in FLT3-KO or control gene-KO HSCs from FL and CB, determined in primary recipients using limiting-dilution assays; 3 to 5 mice per cell dose. **(H)** Hematopoietic lineage distribution based on cell-surface markers expressed in human cells engrafted in mice from (D), determined by flow cytometry. **(I)** NSG-repopulating cell frequencies in FLT3-KO or control gene-KO HSCs from FL and CB, in secondary recipients (sublethally irradiated sex-matched NSG-SGM3 mice), using limiting-dilution assays; 3 to 5 mice per cell dose; human CD45+ cells collected from primary recipients described in (D). Positive engraftment was considered if ≥0.1% human cells; lineage characterization was performed only on grafts with ≥1% of human cells. Mice engrafted with cells with less than 60% KO were excluded from the lineage output analysis. Mice with 0% of human engraftment where no human DNA was detectable were excluded from the genotyping analysis. Unpaired Student’s t test: *P < 0.05; **P < 0.01; ***P < 0.001; ****P < 0.0001; mean ± standard deviation values are reported in the graphs.

To test the importance of FLT3 for human HSC function, we induced CRISPR/Cas9-mediated FLT3-KO in HSCs from FL, CB and HSPCs from BM, followed by xenotransplantation (**Fig 2B**). The percentage of HSC with FLT3-KO pre-transplantation was above 80% in all cell-sources (**Fig 2C**). Human engraftment was successfully generated by FLT3-KO HSCs from FL, CB at 20 weeks and HSPCs from BM at 12 weeks (**Fig 2D**). Compared to controls, engraftment levels were similar in mice transplanted with FLT3-KO cells from CB and BM but significantly lower in mice transplanted with FLT3-KO cells from FL (**Fig 2D** and **Supp.** Fig. 2C). The percentages of cells with KO in xenografts as determined by human-specific PCR were on average above 80% for both FLT3-KO and control groups in FL, CB and BM (**Fig. 2E** and **Supp.** Fig. 2D), establishing that most engrafted cells arose from FLT3-KO HSC. In addition, FLT3 surface expression on phenotypically-defined progenitors at 20 weeks was significantly reduced in the FLT3-KO group compared to control, confirming that the KO was efficacious (**Fig. 2F**).

By limiting dilution analysis (LDA) of 20-week primary grafts, the frequency of repopulating HSCs after KO was similar between FLT3-KO and control groups from FL and CB (**Fig 2G**). Differentiation into all blood-lineages – erythroid, myeloid, T and B lymphoid and megakaryocytic – was not significantly affected by FLT3 KO (**Fig. 2H**). In contrast to the murine system, we did not find differences in the percentages of engrafted human CD34+ cells, nor in the distribution of phenotypic HSCs, multipotent progenitors (MPP) and multi-lymphoid progenitors (MLP) in mice transplanted with FLT3-KO cells from FL and CB compared with controls (**Supp.** Fig 2E). LDA of secondary grafts demonstrated a similar frequency of repopulating cells between FLT3-KO and control groups in CB, but in FL there was a more than 3-fold lower frequency in the FLT3-KO group (**Fig. 2I**). To ensure that our findings were not model-specific, we also used NSGW41 recipient mice that permit human hematopoietic engraftment without the need for irradiation^36^. FLT3-KO did not affect the capacity of FL and CB HSCs to engraft non-irradiated NSGW41 mice (**Supp.** Fig 2F). When we used an alternative pair of gRNAs targeting exon 6 of FLT3 gene (**Supp.** Fig 2G), we obtained results similar to those with our original pair of gRNAs. To provide a positive control for our study, we targeted c-KIT, a gene known to be highly relevant for the HSC function^37–40^. In contrast to FLT3-KO, we observed a drastic reduction in repopulation ability in c-KIT-KO HSC (**Supp Fig 2H**).

Overall, our findings demonstrate that FLT3 is dispensable for the function of normal HSCs from CB and BM, but may be required for maintenance of self-renewal in HSCs from FL.

### 3. FLT3 deletion exposes vulnerabilities specific to ITD+ LSCs and favors normal hematopoietic repopulation

To model the impact of FLT3-KO on a human hematopoietic system composed of normal and leukemic cells, such as found in AML patients, we knocked out FLT3 from CB (CD34+38-) and ITD3 AML (CD34+) cells and co-transplanted them into NSG mice (**Fig 3A**). The percentage of cells with FLT3-KO pre-transplantation averaged 76% in AML and 89.5% in CB (**Supp Fig 3A**), with dominance of non-edited cells in some grafts (**Fig 3B**). For analysis, we divided grafts into those with a high percentage of FLT3-KO cells (FLT3-KO^high^, i.e., > 50% FLT3-KO) and those with a low percentage of FLT3-KO cells (FLT3-KO^low^, i.e., < 50% FLT3-KO), and controls.

**Figure 3.**
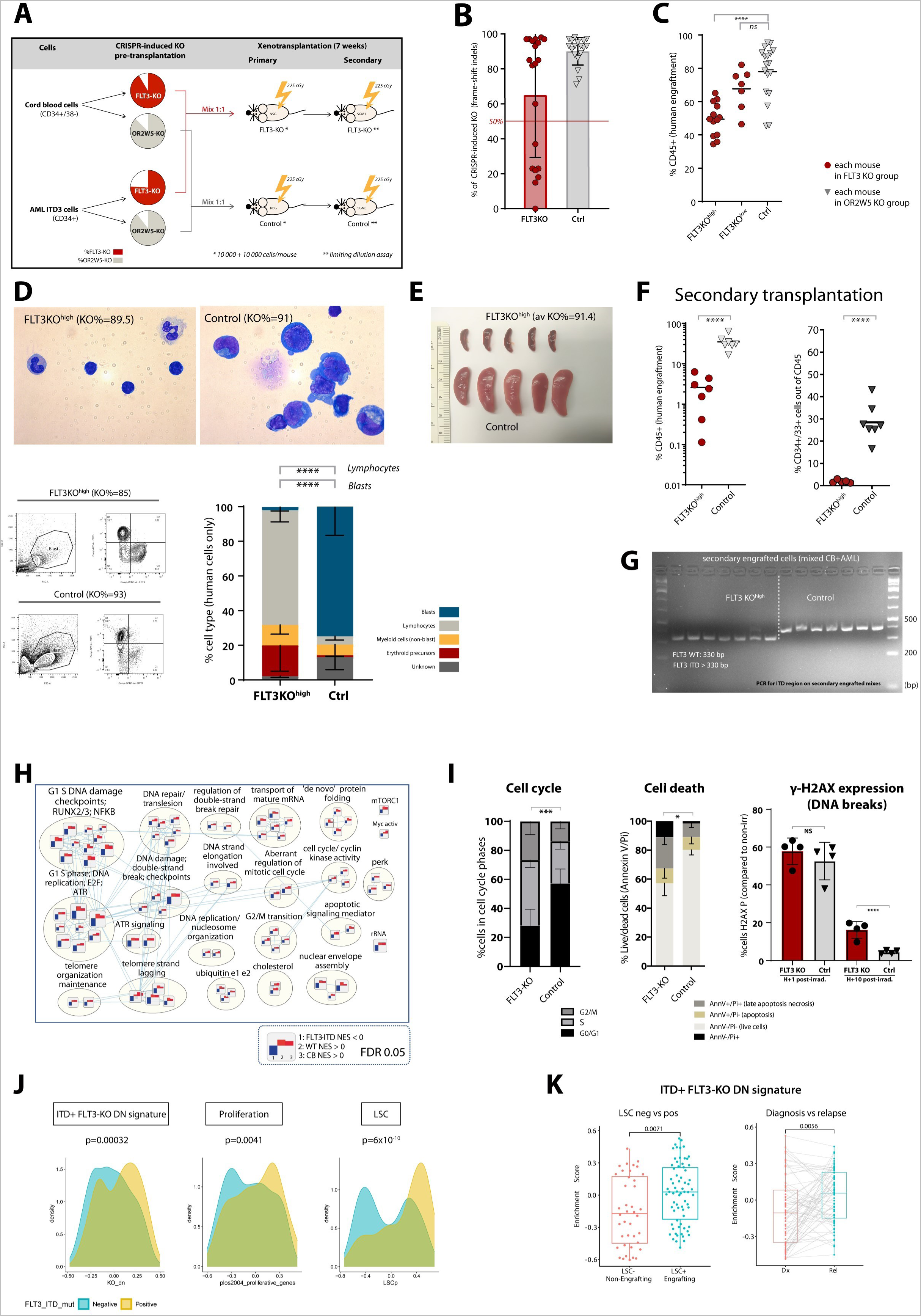
FLT3 deletion disrupt ITD+ LSC-specific pathways and favors normal hematopoietic reconstitution. **(A)** Experimental overview of the competition assay where CB and sample ITD3 were co-transplanted after FLT3-KO and control gene KO. **(B)** KO percentages of human grafts in each mouse transplanted with the mixture (CB and ITD3 cells); 18-20 mice per group; sublethally irradiated NSG female mice; 7-8 weeks of engraftment; experiment performed twice. **(C)** Human engraftment in the competition assay; in the FLT3 KO group, results are divided into grafts with KO percentages > 50% (FLT3KO^high^) and < 50% (FLT3KO^low^); results refer to the injected femur. **(D)** Human cells isolated from the mice bone marrow from (C), prepared by cyto-spinning and stained with Giemsa, 1000X magnification (top); blast gate (SSC/FSC) and expression of myeloid (CD33) and lymphoid (CD19) markers by flow cytometry, representative example (bottom left); quantification of leukemic blasts and normal cells based on cyto-morphology of Giemsa-stained slides from mouse-depleted bone marrow specimens (bottom right), n=3 for each condition; relative to the competition assay. **(E)** Photograph representing spleen size of the mice from the competition assay. **(F)** Secondary transplantation of 125 000 to 1 million human CD45+ cells per mouse, collected from primary FLT3KO^high^ (with KO of ≥ 90%) and control grafts: human engraftment at 7 weeks in sublethally irradiated sex-matched NSG-SGM3 (left); CD34/CD33 expression in human grafts (right). **(G)** Secondary transplantation from (F): assessment of the FLT3-ITD and FLT3-WT alleles in the human engrafted cells in mice from FLT3-KO and control groups, by PCR and gel electrophoresis. **(H)** Gene-pathways downregulated by FLT3-KO specifically in FLT3-ITD AMLs, and not in WT AMLs or normal CB cells; pathway enrichment analysis from the list of the genes differentially expressed by FLT3-KO and control groups of CB (3 samples), ITD-mutated AML (3 samples) and AML without FLT3 mutation (3 samples). **(I)** Cell cycle, cell death and H2AX-P expression by flow cytometry in FLT3-KO and control groups of 2-week grafts from sample AML ITD2; 3-4 sub-lethally irradiated NSG female mice per condition; experimental design in Supp Fig. 3K. **(J)** Expression of the ITD+ *FLT3-KO DN signature* and gene-sets associated with proliferation and LSCs in AMLs with and without FLT3-ITD mutation from 3 independent AML cohorts – TCGA, Leucegene and Beat AML. **(K)** Expression of the *ITD+ FLT3-KO DN signature* in LSC positive (LSC+) and LSC negative (LSC-) cell-fractions from AML samples and in paired diagnosis-relapse samples. Positive engraftment was considered if ≥0.1% human cells; lineage characterization was performed only on grafts with ≥1% of human cells. Unpaired Student’s t test: *P < 0.05; **P < 0.01; ***P < 0.001; ****P < 0.0001; mean ± standard deviation values are reported in the graphs.

FLT3-KO^high^ grafts (**Fig 3C**) were largely composed of normal multilineage populations (**Fig 3D** and **Supp Fig. 3B**) contained <5% blasts (**Fig 3D**), and mice presented normal-sized spleens without blast infiltration (**Fig. 3E** and **Supp Fig. 3C**). By contrast, FLT3-KO^low^ grafts consisted of myeloid-biased leukemic populations (**Supp Fig. 3B**), with bigger spleens partially infiltrated with blasts (**Supp Fig. 3C**). Most of the control group mice were sick, with high levels of engraftment (**Fig 3C**) of a profoundly myeloid-biased leukemic population (**Supp Fig. 3B**), high percentage of blasts (**Fig 3D**) and enlarged spleens massively infiltrated with blasts (**Fig. 3E** and **Supp Fig. 3C**). FLT3-KO^low^ and control grafts contained higher percentages of FLT3+ cells than in the FLT3-KO^high^ group (**Supp Fig. 3D**) and predominantly expressed immature and myeloid markers (**Supp Fig. 3E**) compared to FLT3-KO^high^ grafts where the percentage of cells expressing myeloid/immature markers was drastically reduced and there was an increase in the proportion of cells expressing lymphoid B (CD19) and myeloid mature markers (CD14 and CD15) (**Supp Fig. 3E**). To determine whether leukemic or normal cells dominated engraftment in each mouse, we checked the presence of the ITD versus WT FLT3 allele (**Supp Fig 3F**). Sample ITD3 was dominated by cells with the ITD allele and no WT allele was detected in cells pre-transplant (**Supp Fig 3F**). CB cells were devoid of ITD-mutated DNA, allowing us to infer the presence of cells from AML, CB or both in each graft. Engrafted cells from the FLT3-KO^high^ group only had the WT allele, whereas engrafted cells from the control group mostly had the ITD allele (**Supp Fig 3F**). Secondary recipients engrafted with control cells developed leukemia and exhibited higher levels of human engraftment, higher percentage of CD34+CD33+ immature cells (**Fig. 3F**) and 2-fold higher frequency of repopulating cells (**Supp.** Fig 3G) compared to those engrafted with FLT3KO^high^ cells, who remained healthy and presented normal human grafts (**Fig.3F** and **Supp.** Fig 3G). Upon genotyping, only the ITD allele was detected in control secondary grafts, whereas cells from FLT3-KO^high^ group mostly expressed the WT FLT3 allele (**Fig 3G**). Overall, our results demonstrate that deleting FLT3 abrogates LSC engraftment and self-renewal while sparing normal hematopoiesis.

To investigate the mechanisms underlying the differential effects of FLT3-KO on ITD+ AMLs compared to both normal HSCs and WT AMLs, we performed gene expression analysis (**Supp Fig 3H**). Following FLT3-KO, confirmed at the genomic (**Supp Fig 3I**) and transcriptomic level (**Supp Fig 3J**), cells from ITD+ AML, WT AML and CB samples were cultured short-term and collected for bulk RNA sequencing. Using differentially expressed genes between FLT3-KO and control-edited groups, we performed pathway enrichment analysis and identified the top pathways downregulated (DN) upon FLT3-KO in ITD+ AMLs but not in WT AMLs or CB cells (**Fig 3H**). These were mainly involved in G1 cell cycle phase, G1/S transition in mitosis, cell cycle checkpoints, chromosome maintenance, and DNA damage response (i.e. ATR); together these constitute a signature hereafter called *ITD+ FLT3-KO DN signature*. To evaluate the functional relevance of this signature, we knocked-out FLT3 in sample ITD2, performed 2-week transplants (a time point that allowed detection of FLT3-KO cells) and assayed cell cycle, cell death and DNA repair (**Supp.** Fig. 3K). The proportion of cells in S/G2/M was higher in FLT3-KO grafts than in the control group, whereas the percentage of live cells was lower (**Fig 3I**), indicating that FLT3-KO cells were cycling more frequently, but dying more frequently as well. To test the capacity to repair damaged DNA, 2-week engrafted mice were irradiated 1 hour (h) or 10h before euthanasia, followed by evaluation of γ-H2AX expression in the engrafted cells. At 10h post-irradiation, most control cells had repaired their DNA breaks, whereas 16% of FLT3-KO cells still displayed γ-H2AX expression (**Fig 3I**), indicating that FLT3-KO induced a defect in DNA repair in the context of FLT3-ITD+ leukemia. Expression of the ITD+ FLT3-KO DN signature was upregulated in ITD-mutant AMLs compared to AMLs without ITD mutation from the TCGA, Leucegene and BEAT AML cohorts (**Fig 3J**), validating the relevance of this signature for ITD+ AMLs. In addition, ITD+ AMLs were enriched in proliferative and LSC programs (**Fig 3J**), consistent with previous reports^1,41^. To evaluate the link between our signature and functional LSC, we analysed paired diagnosis-relapse samples and functionally validated LSC+ versus LSC-cell fractions. The ITD+ FLT3-KO DN signature was upregulated at relapse and in LSC+ fractions in all AMLs (**Fig 3K)** as well as in ITD+ mutated AMLs (**Supp Fig 3L).** Overall, our results suggest a link between FLT3, stemness and proliferation in ITD-mutated AML and provide strong evidence that FLT3-KO targets long-term engrafting LSC from FLT3-ITD mutated leukemias through impairment of DNA repair and cell cycle checkpoints.

## DISCUSSION

Here, we have established that the LSCs driving leukemogenesis and relapse in ITD-mutant AMLs have an absolute genetic requirement for FLT3, while it is biologically dispensable for both healthy HSCs and FLT3-WT LSCs. FLT3 has long been a compelling therapeutic target, but after many years of optimization, selective small molecule inhibitors have only demonstrated modest clinical success. By deleting FLT3 on LSCs and HSCs using CRISPR-Cas9 genome editing technology, we have demonstrated that FLT3 targeting is indeed an effective therapeutic strategy as it selectively eradicates LSCs responsible for relapse of FLT3-ITD mutant AMLs without harming HSCs in competitive transplantation. Our work provides a strong rationale for the development of more effective drugs or gene-editing based FLT3 targeting strategies to improve patient outcomes.

Although FLT3 deletion in the mouse causes marked perturbation of lymphopoiesis and other lineages^42^, murine HSC do not express FLT3, making the mouse an unsuitable model system to study the role of FLT3 in human HSCs. Our finding that FLT3 deletion does not impact the ability of neonatal and adult HSCs to reconstitute hematopoiesis provides reassurance of the overall safety of intensifying FLT3 therapeutic targeting in AML. FLT3 deletion did slightly reduce engraftment ability and self-renewal in FL-derived HSCs, however these effects will not likely impact the management of FLT3-mutant AML, which is typically a postnatal disease. FLT3 may play a specialised role in fetal HSCs, which are actively expanding in fetal liver^43,44^ and thus are likely more dependent on growth factors and their receptors including FLT3. In support of this idea, FLT3 is expressed in a transient fetal mouse HSC population that disappears in adult mice^29^.

FLT3-ITD mutations were previously reported to be enriched in defined leukemic stem and progenitor populations using immunophenotypic^45–47^ and transcriptional approaches^1,41^; but the functional effects of FLT3 targeting in LSCs were unclear. The genetic deletion of FLT3 we undertook in this study avoids potentially confounding non-specific effects of pharmacologic inhibition. The demonstration that FLT3-ITD but not FLT3-WT LSCs require FLT3 to form persistent leukemic grafts is intriguing, as leukemic blasts commonly express FLT3 regardless of mutational status^48–50^, as observed in this study. This finding suggests that the reported benefits of FLT3 inhibitors in patients with FLT3-WT AML^51^ may be attributed to multi-kinase inhibition or to inhibition of emerging FLT3-ITD+ clones at relapse rather than to specific effects of FLT3 targeting on WT LSCs.

Consistent with the selective targeting of ITD+ LSCs upon FLT3-KO, our study identified a signature comprising mainly pathways involved in cell cycle checkpoints and DNA repair that were downregulated on FLT3 deletion specifically in ITD+ AMLs. We validated this signature in functional assays, where FLT3-KO increased cell division but also impaired DNA repair, increasing cell death and leading to the extinction of ITD+ leukemic cells. DNA damage and repair has been linked to the FLT3-ITD mutation^52^. Several studies have suggested that some repair pathways work more efficiently in ITD AML than in WT AML, while others are defective^53^. Notably, we demonstrated that our signature is enriched in ITD+ AML from several large independent AML cohorts (TCGA, Leucegene and BEAT AML), confirming its clinical relevance. Cancer stem cells^54^ and specifically LSCs^55^ possess enhanced DNA repair mechanisms that contribute to resistance to genotoxic chemotherapy and radiation. In fact, enrichment of our signature in functionally validated LSCs and at relapse suggests that FLT3 plays a role in DNA repair mechanisms and cell cycle checkpoints in ITD+ LSCs, contributing to the high relapse rates seen in these patients.

Our findings support the design of more potent and specific FLT3 inhibiting drugs, and provide a strong rationale for the development of *in vivo* genetic engineering approaches such as CRISPR/Cas9 genomic engineering^56^ to delete FLT3 in ITD+ AML. In contrast, strategies such as CAR-Ts or BiTEs aimed at eliminating FLT3-expressing cells are not supported by our study, unless special caution is taken to protect HSCs^57^, which express FLT3. The lack of demonstrated effects of FLT3-KO on WT LSCs highlights the need for a multitargeted approach to treat AML, especially considering the genetic heterogeneity and clonal evolution that characterize this disease.

## Acknowledgements

We thank the Orthopedics Department of Centro Hospitalar de São João for assistance with BM collection; the Obstetrics units of Trillium Health, Credit Valley and William Osler Health for CB samples; the Research Centre for Women’s and Infants’ Health Biobank (Mount Sinai Hospital) for FL samples and B. Chow at the Pathology and Laboratory Medicine (Mount Sinai Hospital) for assistance with pathology; M Peixoto for assistance with BM samples and proofreading of the manuscript; M. DSouza and R. Lopez at the Animal Resources Centre (UHN) for support with mouse work; N Simard from from SickKids-UHN Flow and Mass Cytometry Facility and Frances Tong from Princess Margaret Flow Cytometry Facility for assistance with antibody panel design and flow cytometry; Apresto at The Centre for Applied Genomics (SickKids) for sanger sequencing; K. Ho at the Centre for Applied Genomics (SickKids) for next-generation sequencing; K. Asoyan, C. Cimafranca, J. Mouatt, M. Peralta, and Y. Yang at the Pathology Research Program (UHN) and N. Law at the Sttarr Innovation Centre (UHN) for assistance with histology; L. Shultz at the Jackson Laboratory for providing NSGW41 mice; P SCoelho, N Santos, A Zeng, K Kaufman, S Xie for their insight and suggestions; the laboratories of S. Chan and F. Notta for sharing equipment; N. Mbong, S de Silva and M Anders for technical assistance; all members of the J.E.D. Laboratory for critical review of the manuscript.

## Authorship Contributions

J.L.A., J.E.D. and P.P.O. conceived the study. J.L.A and J.E.D..wrote the paper. E.R.L., E.W., O.I.G., J.C.Y.W., P.P.O., M.A.S.S. edited the paper. J.L.A., E.R.L., E.W., O.I.G. analyzed experiments. J.L.A and E.W. designed and performed CRISPR/Cas9 experiments. J.L.A., E.W., E.R.L., O.I.G., B.G. performed *in vitro* and *in vivo* experiments. J.M., L.J., A.M., S.C. assisted with mouse work. J.M. and M.D. performed intrafemoral injections. V.V. and S.B. performed RNA sequencing analysis. A.A. and M.D.M provided primary AML samples, coordinated patient consent and sample collection. J.M.C.C: performed the ITD/WT ratio assay. S.A.performed the smMIP analysis. J.C.Y.W., M.A.S.S. and M.D.M. provided study consultation. J.E.D. secured funding and supervised the study.

## Data availability

Processed transcriptomic data is available at Gene Expression Omnibus (GSE268962) and raw sequence data is in the process of submission to the European Genome-phenome Archive. All other data are available in the manuscript or the supplementary materials.

