## Supplemental data for "FLT3 is genetically essential for ITD-mutated leukemic stem cells but dispensable for human hematopoietic stem cells"

**Supp. Fig 1.**

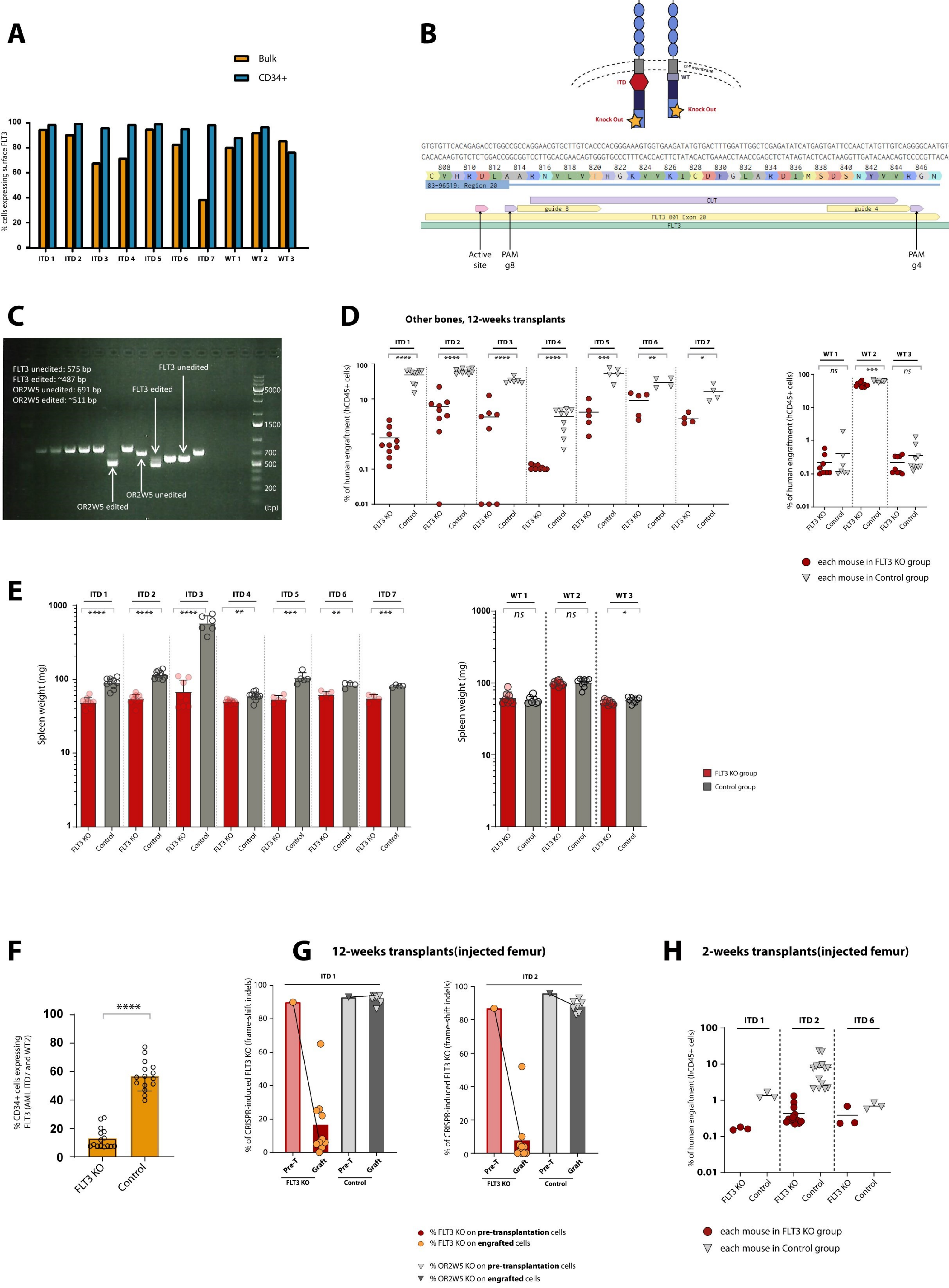
A
Bulk
CD34+
% cells expressing surface FLT3
ITD 1
ITD 2
ITD 3
ITD 4
ITD 5
ITD 6
ITD 7
WT 1
WT 2
WT 3
B
ITD
WT
cell membrane
Knock Out
83-96519: Region 20
CUT
guide 8
guide 4
FLT3-001 Exon 20
FLT3
Active site
PAM g8
PAM g4
C
FLT3 unedited: 575 bp
FLT3 edited: ~487 bp
OR2W5 unedited: 691 bp
OR2W5 edited: ~511 bp
FLT3 unedited
FLT3 edited
OR2W5 unedited
OR2W5 edited
(bp)
D
Other bones, 12-weeks transplants
% of human engraftment (hCD45+ cells)
FLT3 KO
Control
each mouse in FLT3 KO group
each mouse in Control group
E
Spleen weight (mg)
FLT3 KO group
Control group
F
% CD34+ cells expressing FLT3 (AML ITD7 and WT2)
FLT3 KO
Control
G
12-weeks transplants(injected femur)
% of CRISPR-induced FLT3 KO (frame-shift indels)
Pre-T
Graft
% FLT3 KO on pre-transplantation cells
% FLT3 KO on engrafted cells
% OR2W5 KO on pre-transplantation cells
% OR2W5 KO on engrafted cells
H
2-weeks transplants(injected femur)
% of human engraftment (hCD45+ cells)
each mouse in FLT3 KO group
each mouse in Control group

Supp. Fig 2.

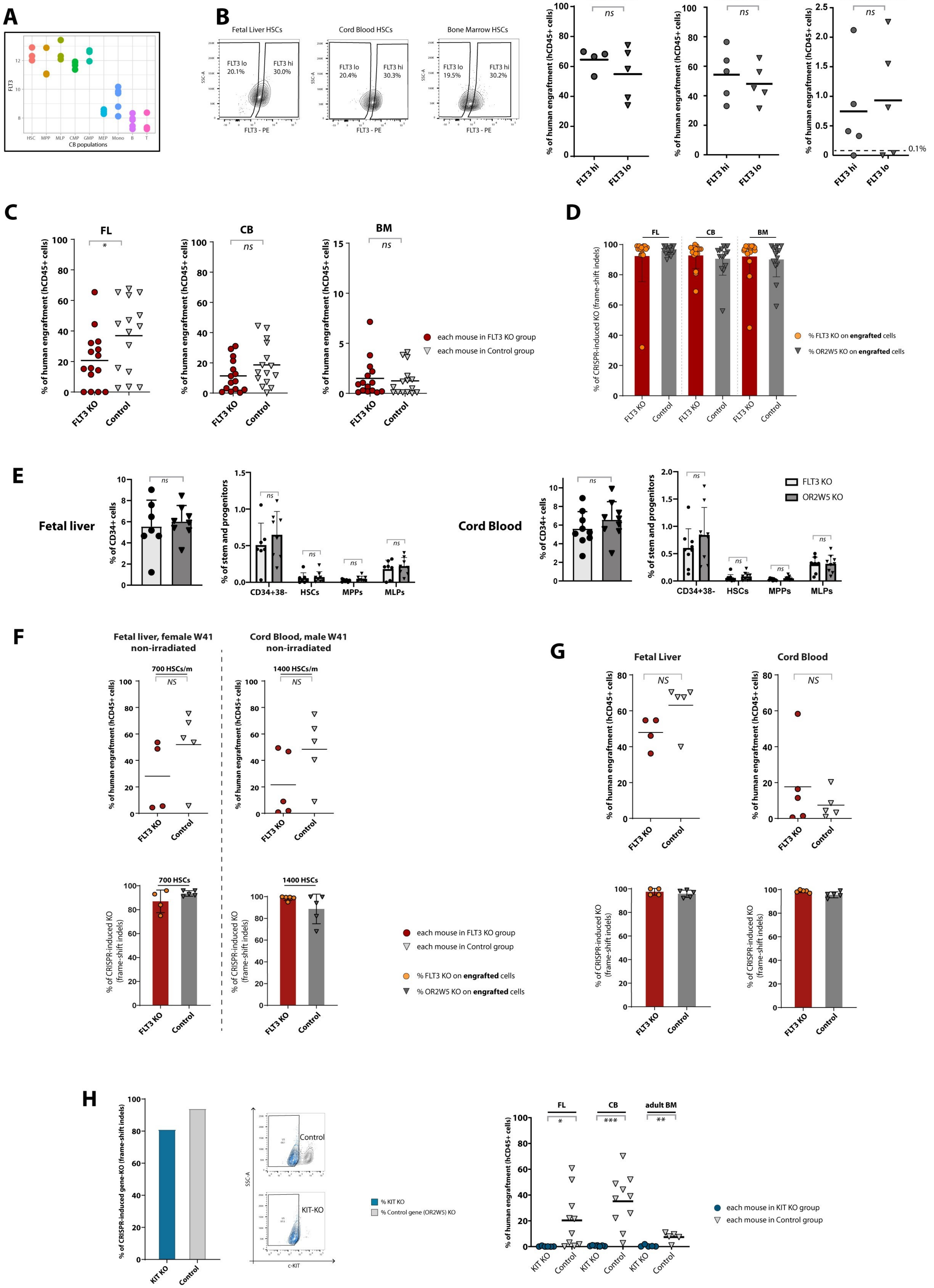
A
B
Fetal Liver HSCs
Cord Blood HSCs
Bone Marrow HSCs
FLT3 lo
FLT3 hi
FLT3 - PE
Fetal Liver
Cord Blood
Bone Marrow
ns
% of human engraftment (hCD45+ cells)
FLT3 hi
FLT3 lo
0.1%
C
FL
CB
BM
*
FLT3 KO
Control
each mouse in FLT3 KO group
each mouse in Control group
D
% of CRISPR-induced KO (frame-shift indels)
% FLT3 KO on engrafted cells
% OR2W5 KO on engrafted cells
E
Fetal liver
Cord Blood
% of CD34+ cells
% of stem and progenitors
CD34+38-
HSCs
MPPs
MLPs
FLT3 KO
OR2W5 KO
F
Fetal liver, female W41 non-irradiated
Cord Blood, male W41 non-irradiated
700 HSCs/m
1400 HSCs/m
NS
700 HSCs
1400 HSCs
G
Fetal Liver
Cord Blood
H
% of CRISPR-induced gene-KO (frame-shift indels)
KIT KO
Control
KIT-KO
SSC-A
c-KIT
% KIT KO
% Control gene (OR2W5) KO
FL
CB
adult BM
***
**
each mouse in KIT KO group
each mouse in Control group

### Supp. figure 3.

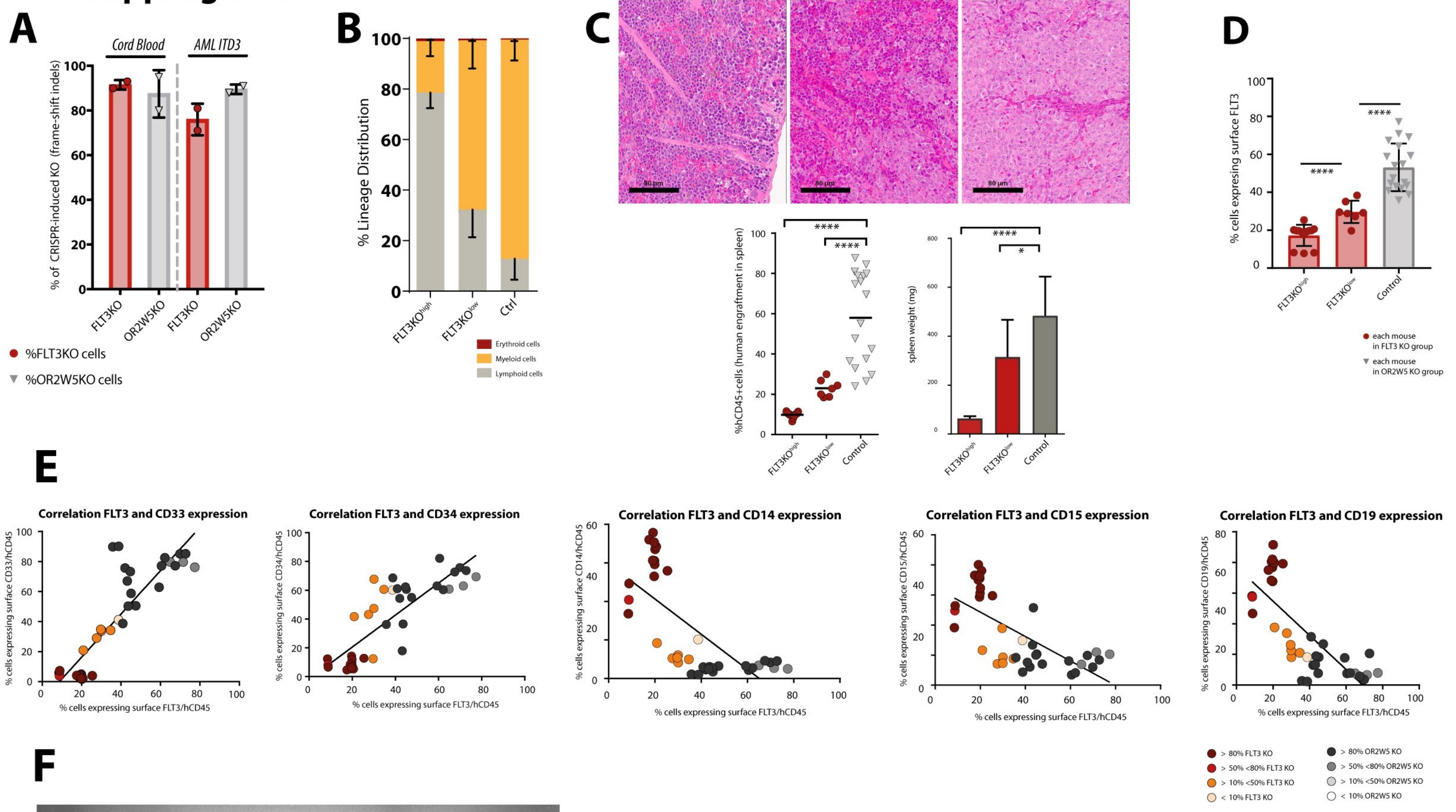


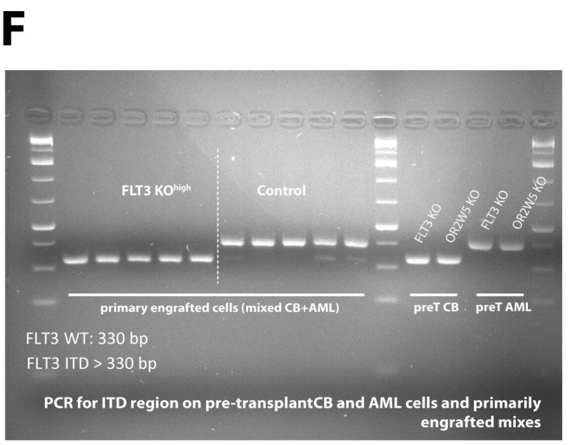


G

| Hematopoietic tissue | Condition | Cell dose hCD45+ | **Secondary** transplant engrafted/injected | Frequency of NSG-SGM3-repopulating cells | *p* |
|---|---|---|---|---|---|
| CB and AML competition | FLT3 KO | 1000000 | 3/3 | 1 in 59 712 | 0.356 |
| | | 125 000 | 4/4 | | |
| | | 15 625 | 0/4 | | |
| | | 1950 | 0/4 | | |
| | OR2W5 KO | 1000 000 | 3/3 | 1 in 27 640 | |
| | | 125 000 | 4/4 | | |
| | | 15 625 | 1/4 | | |
| | | 1950 | 1/4 | | |

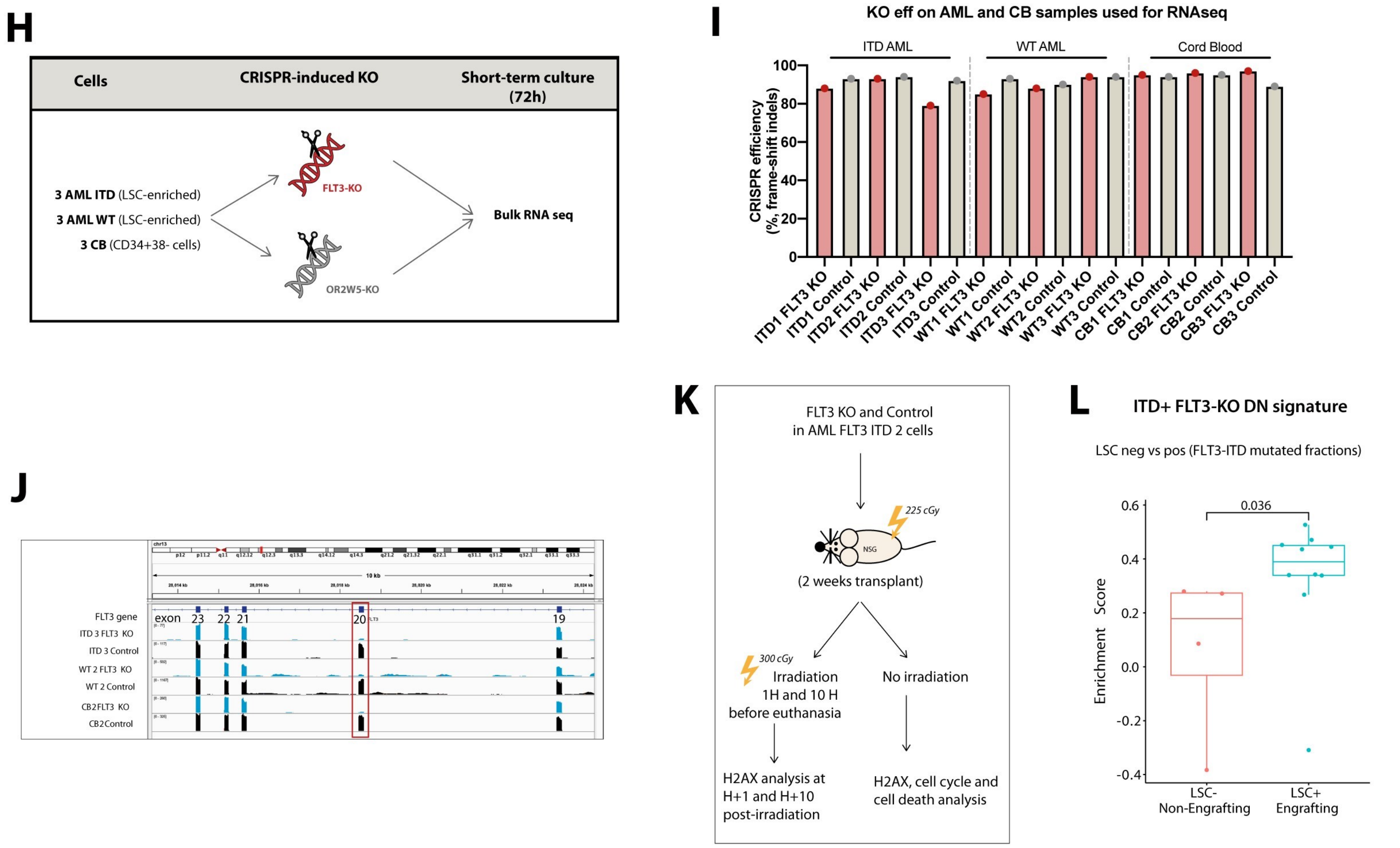

#### Supplemental figure legends

**Supplemental figure 1. Characterization of FLT3-KO in ITD and WT leukemic samples. (A)** Expression of surface FLT3, determined by flow cytometry, on unsorted leukemic blasts and CD34+ sorted leukemic population. **(B)** Top: representation of the FLT3-ITD protein and the WT protein, with the KO site affecting both versions; bottom: representation of the CRISPR/Cas9-edited locus on FLT3 gene, showing the location of the pair of guides, the PAM sequence and the sequence corresponding to the active site of the protein, generated in *benchling.com*. **(C)** Gel electrophoresis analysis of bulk cells cultured for 3 days, derived from CD34+38- FL cells, after FLT3-KO and control gene-KO. **(D)** Human engraftment of leukemic samples at 12 weeks, in sublethally irradiated NSG female mice injected with FLT3-KO or control gene-KO cells; the experiment with sample ITD 7 was humanely terminated at 8 weeks; 4-10 mice per condition; results refer to engraftment in non-injected bones (femur and tibias). **(E)** Spleen sizes of mice from (D). **(F)** FLT3 surface expression of 12-week engrafted cells from samples ITD7 and WT2 - both samples produced grafts with high proportion of FLT3-KO, being possible to test FLT3 expression upon KO; FLT3 expression was determined in the CD34+ population by flow cytometry. **(G)** Samples ITD1 and 2, with FLT3-KO by an alternative pair of gRNA targeting exon 6, in sublethally irradiated NSG mice, at 12 weeks - percentages of FLT3-KO and control gene-KO in the pre-transplantation cells (Pre-T) and in engrafted cells (Graft). **(H)** Short-term (2 weeks) transplants of ITD-mutated AMLs (ITD1, 2 and 6) in sublethally irradiated NSG female mice, human engraftment levels of FLT3-KO and controls; results refer to the injected femur. Positive engraftment was considered if ≥0.1% human cells; lineage characterization was performed only on grafts with ≥1% of human cells. Unpaired Student's t test: *P < 0.05; **P < 0.01; ***P < 0.001; ****P < 0.0001; mean ± standard deviation values are reported in the graphs.

**Supplemental figure 2. FLT3 is expressed in HSCs but it is not required for human engraftment under various conditions. (A)** FLT3 mRNA expression accross human hematopoietic hierarchy, determined by bulk RNA sequencing from 3 independent CB samples. **(B)** FLT3 cell-surface expression in FL, CB and BM HSCs, where FLT3 hi corresponds to the top 30% of cells expressing the highest levels of FLT3, and FLT3 lo corresponds to the bottom 20% of cells expressing the lowest levels of FLT3 (left); human engraftment at 20 weeks in mice injected with 600-800 HSCs FLT3 hi or FLT3 lo from FL, CB and BM, 4-5 mice per group (right). **(C)** Levels of human engraftment in mice injected with FLT3-KO and control gene-KO HSCs from FL and CB and HSPCs from BM, at 20 weeks (FL, CB) and 12 weeks (BM); recipients were sublethally irradiated female NSG mice; results refer to percentages of engraftment in non-injected bones (femur and tibias); 3 human samples of each tissue were used in 3 independent experiments (15 mice per condition); engraftment in injected femurs is represented in Fig. 1D. (**D**) FLT3 and control gene-KO percentages in human cells engrafted in each mouse from (C); KO percentages were determined independently in non-injected bones from each mouse; 1 mouse transplanted with FLT3-KO BM HSCs was not engrafted and human DNA was not detectable so genotyping was not performed. **(E)** Analysis of the stem and progenitor compartment of human engrafted cells in mice transplanted with FLT3-KO and control gene-KO HSCs from FL (right) and CB (left); 2 samples of each tissue, 7-9 mice per condition. **(F)** Human engraftment at 20 weeks, in non-irradiated male and female NSGW41 mice - transplanted with FLT3-KO and control gene-KO HSCs from FL and CB, 4-5 mice per group (top); corresponding FLT3-KO and control gene-KO percentages in the engrafted cells (bottom); number of cells injected per mouse are noted on the top of each graph. **(G)** Human engraftment of HSCs from FL and CB edited for FLT3 with an alternative pair of gRNA targeting exon 6, in sublethally irradiated NSG mice, at 20 weeks, 400-600 HSCs/mouse, 4-5 mice per group (bottom); corresponding percentages FLT3-KO and control gene-KO on the engrafted cells (bottom). **(F)** KIT-KO and control gene-KO percentages in FL cells cultured for 5 days after editing (left); KIT cell-surface expression in the same cells by flow cytometry (middle); human engraftment in sublethally irradiated NSG mice injected with KIT-KO and control gene-KO HSCs from FL and CB and HSPCs from BM, at 20 weeks (FL, CB) and 12 weeks (BM), 5-10 mice per condition (right); for FL and CB experiment, 400-600 HSCs per mouse were injejected; for BM experiment, 3500 HSCs + MPPs were injected; excluded mice engrafted with cells with 0% of KIT-KO (non-edited grafts). Positive engraftment was considered if ≥0.1% human cells; lineage characterization was performed only on grafts with ≥1% of human cells. Unpaired Student's t test: *P < 0.05; **P < 0.01; ***P < 0.001; ****P < 0.0001; mean ± standard deviation values are reported in the graphs.

**Supplemental figure 3. Characterization of FLT3-KO in competition assays and FLT3-KO gene expression signature. (A)** FLT3-KO and control gene-KO percentage in the pre-transplantation CB and AML cells used for the competition assay depicted in Fig 3A; experiment performed twice. **(B)** Hematopoietic lineage distribution based on cell surface markers in human cells engrafted in mice from competition assay depicted in Fig 3A. **(C)** Spleens of the mice from competition assay represented in Fig 3A: hematoxylin eosin staining of paraffin embedded sections of the spleens of xenografts, representative examples (top); human engraftment (bottom left); spleen weight (bottom right). **(D)** FLT3 cell-surface expression in human cells engrafted in mice from the competition assay depicted in Fig 3A. **(E)** Correlation between the expression of primitive, myeloid and lymphoid differentiation markers and FLT3 expression, assessed by flow cytometry; the colors represent the % of gene-KO, each circle represents a mouse from the competition assay. **(F)** Assessment of the ITD and WT allele in FLT3 gene in the pre-transplantation cells (preT CB and preT AML) and in engrafted cells with >50% of FLT3-KO ($FLT3^{high}$) and controls, by PCR and gel electrophoresis, in primary recipients of the competition assay. **(G)** NSG-repopulating cell frequencies in the secondary xenotransplantation assay from Fig 3F, by LDAs. **(H)** Experimental design of the bulk RNA sequencing of FLT3-KO versus control in CB (n=3), ITD-mutated AML (n=3) and AML without FLT3 mutation (WT-AML, n=3). **(I)** FLT3-KO and control gene-KO percentages determined by Sanger sequencing and indel analysis of genomic DNA in the cells used in the experiment from H. **(J)** Representation of the FLT3-KO at the level of the RNA – reduced number of reads of the targeted exon 20 in the FLT3-KO samples compared with control. **(K)** Experimental design of cell cycle, cell death and DNA damage assay of sample AML ITD2 transplanted for 2 weeks in sublethally irradiated female NSG mice. **(L)** Expression of the *ITD+ FLT3-KO DN signature* in LSC+ *versus* LSC- fractions from Fig 3K, but here the analysis was restricted to FLT3-ITD mutated AMLs. Positive engraftment was considered if ≥0.1% human cells; lineage characterization was performed only on grafts with ≥1% of human cells. Unpaired Student's t test: *P < 0.05; **P < 0.01; ***P < 0.001; ****P < 0.0001; mean ± standard deviation values are reported in the graphs.

#### SUPPLEMENTAL METHODS:

##### Human primary samples:

Fetal liver samples were collected from elective pregnancy terminations at 16 to 19 weeks gestation from either sex, with informed consent in accordance with guidelines approved by the Mount Sinai Hospital Research Ethics Board (18-0093-E) and the University Health Network (UHN) Research Ethics Board (02-0763). Samples were processed within 3 hours after collection: each fetal liver was thoroughly sliced using razor blades (VWR) and placed into two 50-mL Falcon tubes previously filled with 40 mL of pre-warmed IMDM medium (Thermo Fisher), 5 mL of collagenase IV (Stem Cell Technologies) and 25 uL of DNAse I at 10 mg/mL (Roche) and incubated at 37° C for 30 min on a shaker. After dissociation, the sample was filtered through a 40 µm cell strainer (Corning), with the help of the black rubber end of a 5 mL syringe (BD). After centrifugation at 350x g for 10 minutes, red blood cells were lysed with 5 mL ammonium chloride (StemCell Technologies) for 5 minutes. IMDM was added to each sample to stop the lysis and the cells were centrifuged at 350x g for 10 minutes. Cell pellets were resuspended in MACS auto running buffer (Miltenyi Biotec) and CD34+ selection was performed using the human CD34 MicroBead kit (Miltenyi Biotec) according to the manufacturer's protocol. CD34+ enriched fetal liver cells were stored in 90% FBS (GE Healthcare) and 10% DMSO (FisherScientific) at -150° C.

Umbilical cord blood samples were collected at Trillium, William Osler and Credit Valley Hospitals with informed consent in accordance with guidelines approved by the University Health Network (UHN) Research Ethics Board. Samples were processed 24-48 h after birth. After dilution 1:1 with phosphate-buffered saline (PBS), mononuclear cells were enriched using lymphocyte separation medium (Multicell). Red blood cell lysis was performed using ammonium chloride solution (StemCell Technologies). For lineage-positive cells depletion, StemSep Human Hematopoietic Progenitor Cell Enrichment Kit (StemCell Technologies) and Anti-Human CD41 TAC (StemCell Technologies) were used for negative selection, according to the manufacturer's protocol. Umbilical cord blood cells, lineage depleted, were stored in 50% PBS, 40% fetal bovine serum (FBS) (ThermoFisher) and 10% DMSO (FisherScientific) at −150 °C.

Bone marrow samples were collected during elective hip replacement surgeries with informed consent and with the approval of the Centro Hospitalar Universitário de São João Ethics Committee (342-18). As part of the hip replacement surgery, femur heads were removed from the patient. Using a bone curette, bone marrow was collected and placed in PBS and processed within 3 hours. Solid pieces of bone and bone marrow were crushed with mortar and pestle and filtered through a 100 µm cell strainer (Corning). Cells were centrifuged at 290x g for 8 minutes and mononuclear cells were isolated using lymphocyte separation medium (Multicell) according to the manufacturer's instructions. After washing, mononuclear bone marrow cells were viably frozen in 50% PBS, 40% FBS (Wisent) and 10% DMSO (FisherScientific) at −150 °C.

Primary AML samples were obtained with written informed consent according to the procedures approved by the University Health Network (UHN) Research Ethics Board (REB 01-0573-C). Samples were isolated as mononuclear cells using lymphocyte separation medium (Multicell) from patients' peripheral blood and frozen viably in FBS (Wisent) plus 10% DMSO.

##### Fetal liver, cord blood and bone marrow cell sorting:

CD34+ fetal liver cells and lineage-depleted cord blood cells were thawed by slowly, dropwise, addition of thawing medium composed of X-VIVO 10 media (Lonza) with 50 % FBS (GE Healthcare) and DNase I (100 µg/ml, Roche). After centrifugation at 350x g for 10 minutes, cells were re-suspended in 1 mL of PBS (Thermo Fisher) and 2.5 % FBS for up to 10 million cells. For bone marrow, an additional previous step of lineage depletion was performed, using StemSep Human Hematopoietic Progenitor Cell Enrichment Kit (StemCell Technologies) and Anti-Human CD41 TAC (StemCell Technologies) for negative selection, according to the manufacturer's protocol. Sorting scheme for hematopoietic stem and progenitor hierarchy protocol was previously described[1]. Briefly, the following antibodies were used: CD45 V500 (1:100, BD), CD34 APC Cy7 (1:200, BD), CD38 PE Cy7 (1:100, BD), CD90 APC (1:50, BD), CD45RA FITC (1:50, BD), CD49f PE Cy5 (1:50). Samples were incubated for 30 minutes at 4° C. After washing, cells were re-suspended in cold 1 mL of PBS (Thermo Fisher) and 2.5 % FBS for up to 10 million cells and Sytox Blue (1:2500, ThermoFisher) was added as a viability die. Cells were sorted on the FACSAria III or FACSAria Fusion (BD). Cell sorting purity checks were performed after each sorting session (accepted purity >95 %). HSCs (CD45+, CD34+, CD38-, CD45RA-, CD90+, CD49f+) and MPPs (CD45+, CD34+, CD38-, CD45RA-, CD90-, CD49f-) were the populations sorted.

**Characterization of AML samples:**

All samples were characterized immunophenotypically by flow cytometry on the FACSAria III or FACSAria Fusion (BD), using the following antibodies: CD45 V500, CD38 BV421 (BioLegend), CD71 FITC, GlyA PE (Beckman Coulter), CD41 PE-Cy5 (Beckman Coulter), CD117 APC, CD34 APC-Cy7, CD15 FITC, CD14 BV605, CD11b PE-Cy5 (Beckman Coulter), CD13 APC, CD33 BV786, FLT3-biotin (1:50) stained with streptavidin PE (1:250) as described below, CD19 V450, CD3 FITC, CD56 BV605, CD7 PE, HLA-DR PE-Cy5 and CD10 APC. All antibodies are from BD unless stated otherwise, and all were used in a 1:100 dilution unless noted differently. Analysis was done on the blast population defined on the SSC/FSC gate.

Samples from patients included in the Princess Margaret Advanced Genomics in Leukemia (AGILE) trial[2] had their mutational composition already characterized by target sequencing. For all the others, we used the Single-molecule molecular inversion probes (smMIPs)-based NGS tools described elsewhere[3].

For the confirmation of the clinical information regarding the presence of the ITD mutation and the absence of the tyrosine kinase domain D835 (TKD) mutation, genomic DNA was isolated using the Agencourt GenFind V3 kit (Beckman Coulter), following the manufacturer's instructions. PCR reaction was composed of 23 µl of genomic DNA, 1 µl of forward and reverse primer (10 µM, IDT) and 25 µl of AmpliTaq Gold 360 master mix (Thermo Fisher). The PCR program was: 95° C for 10 min, followed by 95° C for 30 s, 56° C for 30 s and 72° C for 1 min (40 cycles) and then 72° C for 7 min.

The ITD and TKD assays were adapted from[4]. Briefly, for the ITD assay, the PCR product was analyzed in a 4% agarose gel, to test for the presence of the ITD mutation: 330 bp for the wild-type allele and > 330 bp for the mutant allele.

Primers for the ITD assay:

<u>ITD FWD</u>: 5' GCAATTTAGGTATGAAAGCCAGC 3'

<u>ITD RV</u>: 5' CTTTCAGCATTTTGACGGCAACC 3'

For the TKD assay, the purified PCR product (~8uL) was digested with 1 uL of *Eco*RV (10U/uL, NEB) and 5 uL of restriction buffer REACT2 (Invitrogen) in a final volume of 50 uL (H2O to complete the volume). The mixture was incubated at 37ºC for 2 hours and was subsequently inactivated by heat at 80ºC for 20 minutes. The digested PCR product was analyzed in a 4% agarose gel. The TKD D835 mutation is conveniently located within an *Eco*RV restriction endonuclease digestion site, eliminating this cut site. Therefore, TKD-mutated allele is expected to have 129 bp, wild-type allele 80 bp and undigested fragment 150 bp.

Primers for the TKD assay:

<u>TKD FWD</u>: 5' GTAAAACGACGGCCAGCCGCCAGGAACGTGCTTG 3'

<u>TKD RV</u>: 5' CAGGAAACAGCTATGACGATATCAGCCTCACATTGCCCC 3'

FLT3-ITD allele ratio (FLT3-ITD/FLT3 wild-type) from each ITD-mutated AML sample were evaluated by multiplex PCR. DNA was extracted from patients' peripheral blood mononuclear cells from bulk samples and from CD34+ sorted populations, using a modified protocol of the Agencourt GenFind V2 (Beckman Coulter, A41499) as described previously in this section. PCR amplification was performed using fluorescently labelled PCR primers flanking exons 14 and 15 of FLT3, followed by fragment analysis and allele sizing. FLT3-ITD allelic ratio was determined using the ratio of the peak height of the mutant FLT3-ITD allele over the native FLT3 allele[5]. The limit of detection of this PCR assay is established at 5% for the detection of FLT3-ITD. The primers' sequences used were the same as described earlier in this section for the ITD assay.

**Cell sorting of AML samples:**

AML cells were thawed by slowly, dropwise, addition of thawing medium composed of X-VIVO 10 medium (Lonza) with 50 % FBS (GE Healthcare) and DNase I (100 µg/ml, Roche). After centrifugation at 350x g for 10 minutes, cells were re-suspended in 1 mL of PBS (Thermo Fisher) and 1 % BSA for up to 10 million cells. The following antibodies were used: CD45 V500 (1:100, BD), CD34 APC Cy7 (1:200, BD), CD38 PE Cy7 (1:100, BD), CD3 FITC (1:100, BD). Samples were incubated for 30 minutes at 4° C. After washing, cells were re-suspended in 1 mL of PBS (Thermo Fisher) plus 1% BSA for up to 10 million cells and Sytox Blue (1:2500, ThermoFisher) was added as a viability dye. Cells were sorted on the FACSAria III or FACSAria Fusion (BD). Cell sorting purity checks were performed after each sorting session (accepted purity >95 %). Populations enriched in LSCs were determined on a prior study for samples ITD 1-4 and WT1-3[6]; for samples ITD 5-7, the CD34+ population was selected and the ability to engraft was confirmed *in vivo*. Accordingly, for samples ITD 1 to 7 and WT 1 and 2, CD45+CD34+ population was sorted. For WT3, a sample with low CD34 expression where LSC-enriched population was CD34 negative, the sorted population was CD45+CD3-.

**FLT3 surface expression – cell sorting and analysis:**

For the analysis and sorting accordingly to FLT3 surface expression, the following protocol was designed and optimized for normal and leukemic samples: cells were re-suspended in PBS plus 2.5% FBS or PBS plus 1% BSA as previously described for normal and AML cells, respectively, and stained with FLT3 CD135 biotin-conjugated mouse anti-human CD135, clone 4G8 BD (costume-made, BD, 1:50) for 30 minutes, at 4° C. Cells were washed once with cold PBS plus 2.5% FBS and incubated with streptavidin PE (1:250, BD) for 15 minutes at 4° C, and subsequently washed twice at 4° C. A fluorescent minus one (FMO) control stained with streptavidin PE and the other antibody cocktail excluding FLT3 CD135 biotin was used in all experiments to define the negative gate. For *in vivo* studies, we divided HSC cell population (CD45+, 34+, 38-, 45RA-, 90+, 49f+) by FLT3 surface expression. The top 30% of the population expressing higher levels of FLT3 was considered HSC-FLT3$^{hi}$, whereas the bottom 20% expressing lower levels of FLT3 was considered HSC-FLT3$^{lo}$.

**Guide RNA design:**

Ten guide-RNAs (gRNAs) for FLT3 were predicted using the CRoatan algorithm (http://croatan.hannonlab.org/), and tested in FL, CB, BM and AML cells. From those, we selected a pair of two gRNAs targeting exon 20, near the active site of FLT3 protein (Supp. Fig. 1B), because this pair achieved consistently high percentages of frameshift indels (>80%) at the genomic level (Fig. 2C) which resulted in a drastic reduction of FLT3 expression at the surface of the cells by flow cytometry (Supp Fig1F and Fig. 2F). This pair of gRNAs predominantly caused a deletion of 88 base pairs from exon 20, enabling the detection of the gene-edition by gel electrophoresis analysis (Supp Fig. 1C) - see below "CRISRPR/Cas9 efficiency determination". We also designed eight extra gRNAs on Benchling (http://www.benchling.com). After retesting all eighteen gRNAs for FLT3, we selected another pair targeting exon 6 of FLT3 gene, which were used to confirm the effect of the FLT3-KO in FL, CB and AML.

Four gRNAs for KIT were designed on Benchling and tested in FL, CB and BM cells. We selected one gRNA targeting exon 16, that proved to be efficient at the genomic level (>80% of frameshift indels) and at the protein level, as it caused a significant reduction of KIT surface expression (Supp Fig 2 H).

Control gRNAs were predicted by the CRoatan algorithm to target exon 1 of the olfactory receptor (OR2W5), a gene with no known implications for human hematopoiesis, and the optimization is detailed elsewhere[7]. OR2W5 edition by a pair of gRNAs was detectable by gel electrophoresis analysis (Supp Fig. 1C).

Control gRNA-1: GACAACCAGGAGGACGCACT
Control gRNA-2: CTCCCGGTGTGGACGTCGCA
FLT3 gRNA-4 (exon 20): GATTCCAACTATGTTGTCAG
FLT3 gRNA-8 (exon 20): GGTGACAAGCACGTTCCTGG
FLT3 gRNA-16 (exon 6): GCTTCATGAATTATTTGGGA
FLT3 gRNA-18 (exon 6): TGCCAGAAATGAACTGGGCA
KIT gRNA-4: GCTGAGCTTTTCTTACCAGG

**CRISPR/Cas9 RNP electroporation:**

Sorted HSCs from FL and CB and sorted HSCs combined with MPPs from BM were cultured for 36-48 hours in 96 well round-bottom plates (Corning), in serum-free X- VIVO 10 medium (Lonza) supplemented with 1 % BSA (Roche), 1x L-glutamine (Thermo Fisher), 1x penicillin-streptomycin (Thermo Fisher) containing the subsequent cytokines (Miltenyi Biotec): G-CSF (10 ng/mL), SCF (100 ng/mL), TPO (15 ng/mL) and IL-6 (10 ng/mL). gRNAs were synthesized as Alt-R CRISPR/Cas9 crRNA by IDT, requiring a step of annealing with Alt-R tracrRNA (IDT) to form a functional gRNA duplex. As previously described by us[8,9], CRISPR/Cas9 RNP electroporation was performed using the 4D-Nucleofector (Lonza), chemically synthesized gRNAs (IDT) and recombinant Cas9 nuclease (IDT). Alt-R CRISPR/Cas9 crRNA and tracrRNA (IDT) were re-suspended to 200 μM with TE Buffer (IDT), mixed and heated up to 95° C for 5 min in a thermocycler, then cooled to room temperature at the bench top. In the case of FLT3 and OR2W5, two gRNAs were used to target each respective gene – both crRNAs were annealed to the tracrRNA in a single tube in a 1:1:2 ratio. KIT was only targeted by one gRNA, thus the ratio crRNA/tracerRNA was 1:1. For each electroporation reaction, 1.7 uL of Cas9 protein (IDT), 1.2 uL of crRNA:tracrRNA and 2.1 uL of PBS were combined into a low-binding Eppendorf tube (Axygen, MCT-175-C-S). After 15 minutes of incubation at room temperature, 1 uL of 100 μM electroporation enhancer (IDT) was added. FL, CB or BM cells previously cultured (see above) were washed with warm PBS and centrifuged at 350×g for 10 min at room temperature. Around $10^4$ to $10^5$ cells were then resuspended in 20 uL of Buffer P3 (Lonza) per reaction and rapidly added to the low-binding Eppendorf tube containing the Cas9 gRNA RNP complex. After mixing by pipetting, the mixture was added to the electroporation chamber (Lonza, V4XP3032) and the cells were electroporated using the program DZ-100 in the Lonza 4D-Nucleofector. Quickly after the electroporation, 180 uL of the culture media described above that had been pre-warmed to 37ºC were added to the electroporation chamber. Cells were cultured overnight in 96 well round-bottom plates to recover from the reaction before proceeding to *in vitro* or *in vivo* experiments.

The optimized protocol for primary AML cells required the following alterations: sorted AML cells from each sample were cultured for 24 hours in 24 well flat-bottom plates (Falcon), in serum-free X- VIVO 10 medium (Lonza) with 20% BIT9500 (Stem Cell Technologies), 1% L-glutamine (Thermo Fisher), 1% penicillin-streptomycin (Thermo Fisher) supplemented with IL-6 (20 ng/ml), SCF (100 ng/ml), TPO (25 ng/ml), IL-3 (10 ng/ml), G-CSF (10 ng/ml). The same gRNAs targeting control gene and FLT3 used in normal cells were used, as they provided high editing efficiencies also on primary AML cells. The number of cells per electroporation reaction varied between 2 to 7.5 x $10^6$ and the volumes of the reagents were adjusted accordingly: 8.5 uL of Cas9 protein, 6 uL of crRNA:tracrRNA, 10.5 uL of PBS and 5 uL of 100 uM of electroporation enhancer. Cells were resuspended in 100 uL of Buffer P3 (Lonza) and the electroporation reaction was performed in 100 uL electroporation chambers (Lonza, V4XP-3024) using the program DN-100 in the Lonza 4D-Nucleofector. After the reaction, 1 mL of pre-warmed culture media was immediately added to the electroporation chamber and the cells were cultured overnight to recover before downstream applications. Exceptionally, sample WT3 required the support of stroma cells to be able to survive in the short-term culture required for CRISPR/Cas9 editing. MS5 stroma cells expressing human soluble CSF1 were kindly provided by M.D. Minden's laboratory. MS5 stroma cells were grown in 6-well tissue culture treated plates seeded at a density of 1-1.5 × 10ˆ5 cells/well, and cultured in IMEM medium +10% FBS, 1% Pen/Strep. Cells were propagated in T75 tissue culture treated flasks and passaged every 2-4 days for up to 7-8 passages. Cells from sample WT3 were treated as described for the other samples, except for the utilization of MS5 stroma to support their survival/growth in culture. The rest of the methodology was identical to that used in normal cells.

**CRISRPR/Cas9 efficiency determination:**

A modified protocol of the Agencourt GenFind V2 (Beckman Coulter) was used to isolate genomic DNA. Cells from each condition were transferred to 96-well PCR plates (Eppendorf), enabling the use of multichannel pipetting for each step of the protocol. Into each well, 25 uL of lysis buffer and 1.2 uL of Proteinase K (Zymo Research) were added. After incubation at room temperature for 30 minutes, 50 uL of magnetic particles were pipetted to the well. After 5 min incubation, the plate was placed on a magnetic frame (ThermoFisher) for 10 min. After removing the supernatant with the multichannel pipette, the plate was taken off the magnetic frame and 200 uL of wash buffer 1 were mixed into each well. The PCR plate was placed into the magnetic frame, incubated for 10 min and then the supernatant was removed, and the plate was taken off the magnetic frame. Each well was then washed with 125 uL of wash buffer 2, the plate was placed again onto the magnetic frame and incubated for 10 min, after which the supernatant was removed. The magnetic particles were resuspended with 60 uL of TE buffer (IDT) and the magnetic plate was placed back onto the magnetic frame for 10 min, after which 57 uL of eluted genomic DNA were pipetted into a new PCR plate. PCR was used to amplify each genomic locus edited by CRISPR/Cas9. For each PCR reaction, 23 uL of genomic DNA was mixed with 1 uL of forward and reverse primers (10 uM) and 25 uL of AmpliTaq Gold 360 Master Mix (ThermoFisher). The program in the thermocycler was: 95 °C for 10 min, then 95 °C for 30 s, 56 °C for 30 s and 72 °C for 1 min (x 40 cycles), followed by 72 °C for 7 min. The PCR product was run on a 1.5% agarose gel (ThermoFisher) to check for the presence and the approximate relative amount of edited versus unedited sequences: in controls, a shift in the size of the DNA fragment from 691 to 511 bp was detected from the unedited OR2W5 to the edited fragment; in FLT3-KO, unedited fragments had 575 bp and the edited fragments had 487 bp. As KIT gene was target with only one gRNA it caused small indels and we could not detect a difference between edited or unedited sequences in the gel analysis. In all conditions we proceeded to column purification of the PCR products utilizing the ZR-96 DNA Clean-up Kit (Zymo Research) in accordance with the manufacturer's protocol. The purified PCR product was Sanger sequenced – for OR2W5 and KIT PCR products we used the reverse primer and for FLT3 we used the forward primer. The chromatograms were analyzed using mostly ICE Synthego[10] but also DECODR[11] to characterize CRISPR/Cas9 editing in control-gene, FLT3 and KIT edited DNA. CRISPR/Cas9 efficiency was determined based on the percentage of aberrant sequences after the gRNA cut site. The percentage of CRISPR-induced KO was determined based on the percentage of frameshift indels predicted by the ICE synthego (https://ice.synthego.com) and DECODR (https://decodr.org/) online tools.

PCR primers:
Control (OR2W5) forward primer: 5′-TCGGCCTGGACTGGAGAAAA-3′
Control (OR2W5) reverse primer: 5′-GAGACCACTGTGAGGTGAGA-3′
FLT3_exon20 forward primer: 5′- CAGAAGCCGCACAAAGAACTG -3′
FLT3_exon20 reverse primer: 5′- CTAGCTAGCCCAAGGACAGATG -3′
FLT3_exon6 forward primer: 5′- CTCTGAGCTCTTAGCTGAAACCA -3′
FLT3_exon6 reverse primer: 5′- TGCATTTACTGTGACCTGAAGGA -3′
KIT forward primer: 5′- TAGGGTTGCAAGTGGGTGTTT -3′
KIT reverse primer: 5′- GTCTATGCTTCACCCCAAGCC -3

**Animal studies:**

All mouse experiments were approved by the Animal Care Committee of the University Health Network (UHN) and were performed in accordance with all the relevant regulatory and ethical standards. Primary xenotransplantation experiments were performed in 8- to 12- week-old female *NOD.Cg-PrkdcscidIl2rgtm1Wjl/SzJ* (NSG) mice (JAX), whereas secondary xenotransplantation experiments were performed in 10- to 16- week-old sex-matched NOD.Cg-Prkdc$^{scid}$Il2rg$^{tm1Wjl}$Tg(CMV-IL3,CSF2,KITLG)1Eav/MloySzJ (NSG-SGM3) mice (JAX), bred at the University Health Network/Princess Margaret Cancer Center. Both NSG and NSG-SGM3 were sublethally irradiated with 225 cGy, 24 hours before transplantation. We also used 8- to 12-week-old female and male NOD.Cg-PrkdcscidIl2rgtm1WjlKitem1Mvw/SzJ (NSGW41) mice, a kind gift from Leonard D. Shultz (The Jackson Laboratory). NSGW41 mice can support human hematopoiesis without the need for irradiation, providing an opportunity to test the effect of FLT3-KO on the ability of human hematopoietic cells to engraft non-irradiated recipients.

**Xenotransplantation in primary recipients:**

For xenotransplantation of FL and CB cells, at least 400 HSCs (CD45+, CD34+, CD38-, CD45RA-, CD90+, CD49f+) per mouse were used; for BM experiments, at least 800 to 1000 HSCs +/- 800 to1000 MPPs (CD45+, CD34+, CD38-, CD45RA-, CD90-, CD49f-) were used per mouse; for AML, the cell dose was calculated based on previous LDAs performed in our laboratory[6] for each individual AML sample. Those cell-doses for each different type of sample – FL, CB, BM and AML – were defined to achieve a robust engraftment. For each experiment, the same cell dose was used in all mice (experimental group and control). Please, see Supp. Table 1 for detailed information about the main experiments – FLT3, KIT and control-gene KO in FL, CB, BM and AML cells transplanted into NSG mice. The cell dose and other details on the confirmatory experiments using FLT3$^{high}$ versus FLT3$^{low}$ HSCs, alternative guides for FLT3 KO, and alternative recipients (NSGW41 mice) are noted on the respective figures and figure legends.

For FLT3$^{high}$ versus FLT3$^{low}$ experiment, cells were injected into mice directly after FACs-based cell sorting. For all experiments where CRISPR-editing was performed, cells were sorted, cultured for 48 hours (normal cells) or 24 hours (AML cells), electroporated with CRISPR/Cas9 machinery and cultured over-night, before injection. Exceptionally, sample WT3 required the support of stroma cells to be able to survive in the short-term culture required for CRISPR-Cas9 editing – please, see "CRISPR/Cas9 RNP electroporation" for further details.

Xenotransplantations were performed by intrafemoral injections, as described in[12]. Briefly, mice were anesthetized with isoflurane and the right femur was perforated through the knee with a 27-gauge needle. Human cells, diluted in 30 uL PBS, were then injected into the femoral cavity through a 28-gauge ½ cc syringe (Becton Dickinson). Mice transplanted with FL and CB cells were euthanized at week 20, whereas mice transplanted with BM cells were euthanized at week 12, to ensure that the grafts were still detectable, except if stated otherwise. Mice transplanted with leukemic cells were euthanized at week 12 or whenever human endpoints were reached, as noted in the results section. The injected bone (femur) and the other bones (tibias, and non-injected femur) were flushed with 1 ml PBS + 2.5 % FBS. Spleens were collected and crushed using the end of the plunger of a 5 mL syringe (BD) and pushed through a 5 mL tube with a 35 uM cell strainer (Corning). Cells were centrifuged at 350xg for 10 min and re-suspended in 500 uL of PBS + 2.5% FBS and counted in 500 µl ammonium chloride (Stem Cell Technologies) using the Vicell XR (Beckman Coulter). For subsequent genotyping, 25 uL of cells were to a PCR plate (Eppendorf) and stored at -80° C.

For flow cytometry analysis, 50 uL of cells were mixed with 50 uL of antibody mix for 60 minutes at 4ºC for staining. We used the following antibodies (all form BD unless stated otherwise) to characterize the surface expression of lineage markers in all experiments: CD45 AF700 (1:100), CD19 V450 (1:100), CD3 FITC (1:100), CD33 APC (1:100), CD41 PE-Cy5 (1:200, Beckman Coulter), GlyA PE (1:100, Beckman Coulter), CD71 BV650 (1:100) and CD34 APC-Cy7 (1:100). For AML experiments, the following antibodies were additionally used to characterize blast surface expression: CD45 V500, CD38 BV421 (BioLegend), CD34 APC-Cy7, FLT3-biotin (1:50) stained with streptavidin PE (1:250) as described, CD33 BV786, CD15 FITC, CD14 BV605, CD11b PE-Cy5 (Beckman Coulter), CD13 APC, CD7 PE, HLA-DR PE-Cy5 and CD117 APC (all antibodies from BD and all 1:100 unless stated otherwise). Cells were analyzed on the FACSCelesta (BD) with a high throughput sampler (HTS). Cells from each mouse (injected femur and other bones, separately) were viably frozen and stored at -150° C. All flow cytometry analysis was performed using FlowJo (BD) in a blinded manner. Only mice with ≥ 0.1% of CD45+ engraftment level in the injected femur or in the other bones were considered engrafted. For lineage distribution in normal cells and for blast characterization in leukemia, only mice with ≥ 1% of CD45+ engraftment were used, because characterization of sub-populations with lower levels of engraftment would risk being inaccurate.

**Supplemental Table 1. Methodological details of the main *in vivo* experiments - FLT3-KO, KIT-KO and OR2W5-KO in FL, CB, BM and AML cells transplanted into sublethally irradiated NSG mice.**

| | Experimental condition | Human tissue (n) | Cells/mouse | Duration |
|---|---|---|---|---|
| Normal cells | FLT3 -KO *versus* control | Fetal liver (3) | ~600 HSCs | 20 weeks |
| | | Cord blood (3) | ~400-600 HSCs | 20 weeks |
| | | Bone marrow (3) | ~2000-4500 HSCs+MPPs | 12 weeks |
| | KIT-KO *versus* control | Fetal liver (2) | ~600 HSCs | 20 weeks |
| | | Cord blood (2) | ~400 HSCs | 20 weeks |
| | | Bone marrow (1) | ~3500 HSCs+MPPs | 12 weeks |
| Leukemia | FLT3 -KO *versus* control | ITD 1 | ~1000 000 CD34+ | 12 weeks |
| | | ITD 2 | ~400 000 CD34+ | 12 weeks |
| | | ITD 3 | ~20 000 CD34+ | 8 weeks* |
| | | ITD 4 | ~400 000 CD34+ | 12 weeks |
| | | ITD 5 | ~700 000 CD34+ | 12 weeks |
| | | ITD 6 | ~900 000 CD34+ | 12 weeks |
| | | ITD 7 | ~160 000 CD34+ | 12 weeks |
| | | WT 1 | ~150 000 CD34+ | 12 weeks |
| | | WT 2 | ~100 000 CD34+ | 12 weeks |
| | | WT 3 | ~1000 000 CD45+CD3- | 12 weeks |

* Animals transplanted with sample ITD3 were humanely euthanized at 8 weeks.

**CRISPR/Cas9 editing on *in vivo* assays:**

After CRISPR/Cas9 RNP electroporation, most cells were transplanted into mice, but a small portion was cultured for 3-5 days, with the respective medium for normal and leukemic cells (see above). From these cells, genomic DNA was isolated, the CRISPR/Cas9 edited locus was amplified by PCR, and Sanger sequencing was performed as described above. Chromatograms were analyzed using mostly the online tool ICE Synthego, but also DECODR, to determine the percentage of edited DNA in the cells from both FLT3-KO and control gene-KO conditions, thus determining the *pre-transplantation KO percentage*. To determine the *post-transplantation KO percentage* in each xenograft, genomic DNA was isolated form the bulk bone marrow cells of the injected bones and other bones individually, and Sanger sequencing and chromatogram analysis on ICE synthego platform were carried out as described. All PCR primers were human specific. Please, see the section "CRISRPR/Cas9 efficiency determination" for further details.

In all experiments, as mentioned above, engraftment was considered positive when hCD45 was ≥ 0.1% in the bone marrow and lineage output was only analyzed when human grafts were ≥ 1%.

In the normal HSPCs experiments, for which we used FL, CB and BM cells, all samples were included in the % of human engraftment and in the determination of KO percentage, except for mice with 0% of human engraftment where no human DNA was detectable, which were excluded from the genotyping analysis. For the lineage output, only mice with KO percentages above 60% were included, in order to conclude about the effect of the KO on the ability to differentiate. These criteria were defined in advance of analysis.

In the leukemic experiments, all samples were included in the % of human engraftment and in the determination of KO percentage. Regarding the FLT3-KO percentage results of the *post-transplantation* samples from the ITD group (Fig 1), it was very important to ensure that the grafts were composed of human leukemic cells, even when KO percentages were low. Therefore, all engrafted samples were included in the lineage output analyses.

Confirmation of human specificity of all primers used was done by performing PCRs in murine cells, and verifying that DNA was not amplified in the absence of human cells. Whenever the percentage of KO is referred to be very low or zero, it applies to successfully amplified human DNA that does not show any CRISPR/Cas9-mediated editing.

**ITD ratio determination on ITD-mutated AML engrafted samples:**

ITD allele ratio was determined for the ITD-mutated samples engrafted on primary xenotransplants. Bone marrow from each mouse engrafted with ITD-mutated AML samples was thawed and depleted from murine cells using the mouse cell depletion kit (Miltenyi Biotec). A modified protocol of the Agencourt GenFind V2 (Beckman Coulter) was used to isolate genomic DNA, as described in "CRISRPR/Cas9 efficiency determination". The ITD locus was amplified by PCR as described in the ITD assay in the section "Characterization of AML samples". ITD ratio for each sample was determined as described in "Characterization of AML samples".

**Short term transplantation**

For short-term transplantation of leukemic samples, a larger number of cells was needed to obtain measurable human engraftment at 2 weeks, limiting this experiment to those samples with high numbers of cells. The procedure was performed as described for the 12-weeks transplant, except for the number of cells: for the sample ITD1, 2.5 million cells were injected per mouse, with 3 mice per condition and the experiment was done once; for the sample ITD2, 1.6 to 3 million cells were injected per mouse and the experiment was repeated 3 times, using a total of 14 mice per condition; For the sample ITD6, 3.5 million cells were used per mouse, with 3 mice per condition, and the experiment was performed once. Cells were injected in sublethally irradiated (225 cGy) NSG mice, by intrafemoral injections as described, and the experiment was terminated at 2-weeks. Human engraftment analysis and CRISPR-Cas9 editing determination were performed as described for the other primary xenotransplantation assays.

**Xenotransplantation in secondary recipients and limiting-dilution assays:**

Frozen samples of each mouse of primary xenotransplantations, previously checked for >80% CRISPR-mediated KO efficiency, were individually thawed. All the samples from each mouse (injected bones and the other bones) were combined and mouse-cell depletion was performed (each mouse individually), using the mouse cell depletion kit (Miltenyi Biotec) - cells were stained for 15 min at 4° C and then transferred to LS columns (Miltenyi Biotec). Samples from each mouse were analyzed individually using stem and progenitor markers: : CD45 V500 (1:100, BD), CD34 APC Cy7 (1:200, BD), CD38 PE Cy7 (1:100, BD), CD90 APC (1:50, BD), CD45RA FITC (1:50, BD), CD49f PE Cy5 (1:50) and FLT3 CD135 biotin-conjugated antibody (1:50, BD, costume-made) followed by secondary staining with streptavidin PE (1:250, BD), as described above. For this, up to $1x10^6$ cells were re-suspended in 500 uL of PBS with 2.5% FBS, stained with all antibodies except for streptavidin for 30 minutes at 4ºC, washed and then stained with streptavidin-PE for 15 minutes at 4ºC and washed twice. Samples were individually analyzed on the FACSAria III or FACSAria Fusion (BD), live cells were sorted and pooled according to each condition – FLT3 KO and control - using 6 to 8 mice per condition from each experiment. For the limiting-dilution assays, pooled cells from each condition were transplanted at defined cells doses into secondary recipients – NSG-SGM3 – via intrafemoral injections, as described. Eight to twelve-weeks later, mice were euthanized and bone marrow from the injected femur and the contralateral femur were collected separately and analyzed. Engraftment was scored as positive if the percentage of human CD45+ cells was ≥ 0.1% in the bone marrow. The frequencies of NSG-repopulating cells in secondary recipients were estimated by linear regression analysis and Poisson statistics using the publicly available online tool ELDA[13] (http://bioinf.wehi.edu.au/software/elda/). Human grafts were analyzed by flow cytometry as described for primary xenotransplantation assays.

To determine the frequencies of NSG-repopulating cells in primary recipients by LDA, FL and CB HSCs were sorted, CRISPR-Cas9 edited as described and transplanted at defined cell doses (as indicated in the Fig. 1E) into primary NSG mice. The analysis was performed as described for secondary xenotransplantation experiments.

**Competition assay of CB and ITD-mutated AML (primary and secondary)**

CD34+38- cells were sorted from cord blood and CD34+ cells were sorted from the leukemic sample ITD3, as described in the cell sorting sections. FLT3-KO and control gene-KO were performed in both samples individually, as described in CRISPR/Cas9 RNP electroporation. CD34+38- CB cells and CD34+ ITD3 populations contain a similar frequency of long-term repopulating cells allowing for a competitive repopulation experiment with a 1:1 proportion of cells (10 000 to 20 000 cells from each per mouse). Prior to transplantation, CB cells were mixed with ITD3 cells in a 1:1 proportion and transplanted by intrafemoral injection into sub-lethally irradiated (225 cGy) female NSG mice. Ten mice received a mixture of CB and ITD3 cells with FLT3-KO, and the other ten mice received the mixture with OR2W5-KO. This experiment was performed twice (total of 20 mice per group). Mice were euthanized at 7 to 8 weeks, respecting humane endpoints. Human engraftment, lineage output, blast panel analysis and CRISPR/Cas9-mediated KO percentage were determined as described in “xenotransplantation in primary recipients” and “CRISPR/Cas9 editing on in vivo assays” sections. For cyto-morphology analysis, pooled samples from each group were depleted from mouse cells, human cells were cyto-spun and stained with Giemsa (n=3 pools for each condition), as detailed in the section “Cyto- and histo-morphology” below.

For secondary transplantation, samples engrafted in primary recipients with > 90% of CRISPR/Cas9-mediated gene-KO were selected from both FLT3-KO and control groups, thawed and mouse-cell depletion was performed as described in the “Xenotransplantation in secondary recipients and limiting-dilution assays” section. Samples were individually analysed using stem and progenitor markers as described, and live/human CD45+ cells were sorted, pooling samples from 6 to 7 primary mice from each condition (FLT3-KO and control). For the limiting-dilution assays, pooled cells from each condition were transplanted at defined cells doses into secondary recipients – NSG-SGM3 – via intrafemoral injections, as described.

The presence of the ITD mutation was tested both in CD34+38- CB cells and CD34+ ITD3 cells, separately, and in the engrafted cells in primary and secondary recipients, as described in “Characterization of AML samples”. The presence of the ITD allele, the WT allele or both was used as an indication of which competing cell-population engrafted – CB, leukemia or both.

**DNA damage, cell cycle and apoptosis analysis *ex vivo***

Short-term transplants were performed using the sample ITD2, as described in "Short term transplantation". After FLT3-KO and control gene-KO, 3 million cells per mouse from ITD2 sample were transplanted into sublethally irradiated female NSG-mice, by intrafemoral injections. After 2 weeks, FLT3-KO and control mice were divided into 3 groups: 1 group was irradiated with 300 cGy and euthanised an hour later (4 mice per condition), another group was irradiated with the same dose but euthanised 10 hours later (4 mice per condition) and a group of mice was not irradiated (3 mice per condition). Bone marrow from the injected bones and the other bones was flushed with 1 ml PBS + 2.5 % FBS, as described above. Subsequently, 50 uL of each flushed bone marrow were collected for genotyping of the CRISPR/Cas9 edits and 150 uL were used for staining with hCD45 AF700 (1:100, BD) and Annexin V/Propidium iodide (Pi) using BD Annexin V: FITC Apoptosis Detection Kit II, following manufacturer's instructions.

The remaining volume flushed from each bone was pooled into one tube per mouse and murine cells were depleted using the mouse cell depletion kit (Miltenyi Biotec) as described before. Cells were mixed with 4% PFA pre-warmed to 37ºC (10 volumes) and incubated at 37ºC for 12 minutes. After washing, cells were re-suspended in PBS with 2% FCS and mixed with PermBuffer III (BD Phosflow) directly from -20ºC freezer, and the mixture was incubated on ice for 30 minutes. After washing twice, cells were re-suspended in PBS with 2% FBS and blocked with FcR Blocking Reagent, human (1:100, Miltenyi Biotec). Anti-H2AX (pS139) PE (1: 100, BD Pharmigen) was added, and the mixture was incubated for 1 hour on ice in the dark. After washing, cells were re-suspended in PBS with 2% FBS, Pi (2 ug/mL, BD) and RNase A (10 ug/mL, Ambion) were added. Samples were analyzed on the FACSCelesta (BD).

All flow cytometry analysis was performed using FlowJo (BD) in a blinded manner. Staining for cell cycle and H2AX-P was performed in mouse-depleted cells to ensure only human cells were being analyzed. With the same goal, cell death/apoptosis was analyzed on the hCD45+ population, as mouse-cell depletion may create important artifacts on this analysis. The analysis for cell cycle and apoptosis was performed on mice that did not receive irradiation before euthanasia, whereas for the DNA damage and repair analysis, both mice that did not receive and that receive irradiation before euthanasia were used. Human engraftment and CRISPR/Cas9-mediated KO percentage were determined as described in "xenotransplantation in primary recipients" and "CRISPR/Cas9 editing on in vivo assays" sections.

**Cyto- and histo-morphology**

In xenotransplantation assays, multiple samples of flushed bone marrow of engrafted mice were combined and mouse-depletion was performed using the mouse cell depletion kit (Miltenyi Biotec). After staining for 15 min at 4° C, cells were transferred to an LS column (Miltenyi Biotec). Collected human cells were placed into Shandon cytofunnels (Thermo Fisher) and spun using the CytoSpin 4 instrument (Thermo Fisher) at 112x g for 10 min with medium acceleration. After air-drying overnight, slides were stained using a standard Giemsa staining (Sigma). Mice spleens were collected and fixed in 10 % formalin solution (Sigma) for 24 hours at 4ºC and subsequently embedded in paraffin, sectioned and stained with a hematoxylin and eosin protocol; stained slides were scanned on the AT2 slide scanner (Aperio ImageScope, Leica). All cell observations and quantifications were performed in a blinded manner.

**RNAseq:**

Library preparation and sequencing

Cells from AML samples ITD 1, ITD 2, ITD 3, WT 1, WT 2 and WT 3 were thawed and the CD34+ population was sorted from all samples, except from WT 3 where the CD45+ CD3- population was sorted, as described in "Cell sorting of AML samples". Three CB samples were thawed and the CD34+CD38- population was sorted as described in "Fetal liver, cord blood and bone marrow cell sorting". AML samples were cultured for 24-h, while CB cells were cultured for 48-h, using the optimal culture conditions for each type of sample. Subsequently, FLT3-KO and OR2W5-KO were performed using CRISPR/Cas9 technology through electroporation – culture conditions and

CRISPR-editing detailed in "CRISPR/Cas9 RNP electroporation". After electroporation, cells were cultured for 3 additional days to achieve the maximal reduction on FLT3 expression after FLT3-KO in culture. Subsequently, cells were collected and viably sorted using SYTOX Blue Dead Cell Stain (Thermo Fisher Scientific). A minimum number of 13 x$10^3$ live cells was sorted per sample (range 13 to 800 x$10^3$). One fifth of the sample was collected for total DNA isolation and subsequent determination of KO percentage mediated by CRISPR/Cas9 (as described in "CRISRPR/Cas9 efficiency determination"). The rest of the sample was used for total RNA purification and DNase treatment using the RNeasy Micro kit (Qiagen). RNA integrity was measured using the RNA 6000 Pico kit (Agilent) and the average RNA integrity scores were the following: 8 for AML-ITD samples (6.9-10), 10 for AML-WT samples (9.7-10) and 9.95 for CB samples (9.8-10). To generate RNAseq libraries, total RNA was used as input with SMART-Seq V4 Ultra Low Input RNA kit (Clontech). The cDNA profiles were confirmed on the bioanalyzer using the High Sensitivity 14 DNA kit (Agilent) and next-generation sequencing libraries were made using the Nextera XT DNA Library Preparation kit (Illumina). Sequencing was performed on the NovaSeq 6000 system using 2 x 150 bp paired-ended sequencing to obtain ~50 M reads per sample.

Alignment and Differential expression

Reads were aligned with STAR v2.5.2b[14] against hg38 and annotated with ensembl v90. Default parameters were used except for the following: chimSegmentMin 12; chimJunctionOverhangMin 12; alignSJDBoverhangMin 10; alignMatesGapMax 100000; alignIntronMax 100000; chimSegmentReadGapMax parameter 3; alignSJstitchMismatchNmax 5 1 5 5. Read counts were generated using HTSeq v0.7.2[15]. Differential gene expression was performed using edgeR_4.0.16[16] in R 4.3.2 following recommended practices.

Pathway enrichment analysis

Pathway enrichment analysis and visualization was performed as described previously[17]. Briefly, a score to rank genes from top upregulated to downregulated was calculated using the formula -sign(logFC) * -log10(pvalue). The rank file from each comparison between FLT3 ITD KO and corresponding controls was used in GSEA analysis (https://www.gsea-msigdb.org/gsea/index.jsp) using 2000 permutations and default parameters against indicated gene sets. All gene sets were obtained from a pathway database (http://baderlab.org/GeneSets. EnrichmentMap version 3.4.0) and AutoAnnotate 1.5.0 in Cytoscape 3.10.2 were used to visualize enriched gene-sets with indicated FDR-q value and NES and a Jaccard coefficient set to 0.375

GSVA on public AML data

We applied GSVA with "Gaussian" kcdf to transcripts per kilobase million (TPM)-normalised gene expression from the TCGA, BEAT-AML and Leucegene AML cohorts to generate ITD+ FLT3-KO down signature-specific enrichment scores for each patient as well as other signatures: "Proliferation"[18], and "LSC"[6]. We applied GSVA to TPM-normalised RNA-sequencing from primary AML cell-sorted fractions evaluated for functional LSC activity via xenotransplantation[19]. Log(TPM) expression was compared between LSC+ and LSC- fractions using a two-tailed unpaired t test. We also filtered our dataset for 14 LSC samples with a positive FLT3-ITD mutation to assess whether our signature was enriched in LSC+ fractions for all AML samples and specifically FLT3-ITD-mutated samples. To determine whether our signature was associated with relapse, we analysed the log(TPM) expression of 68 paired AML samples collected at diagnosis and relapse following chemotherapy, compiled from 5 independent cohorts, using paired t tests based on the significance of their enrichment at relapse[20–23].

**Statistical analysis:**

GraphPad Prism v9.0 or Microsoft Excel were used for statistical analysis, except for RNA-sequencing (analysis detailed in the corresponding section). Statistical significance was calculated using two-tailored unpaired student's T-test for most experiments. One-way ANOVA was applied to compare means between three or more independent groups. Mean ± standard deviation values are reported in the graphs. Sample size was selected to provide sufficient power for calling significance with standard statistical tests.

| REAGENT or RESOURCE | SOURCE | IDENTIFIER |
|---|---|---|
| **Antibodies** | | |
| Mouse monoclonal anti-CD3, FITC, clone SK7 | BD | 349201; RRID:AB_400405 |
| Mouse monoclonal anti-CD7, PE, clone M-T701 | BD | 340581; RRID:AB_400064 |
| Mouse monoclonal anti-CD10, APC, clone HI10a | BD | 340923; RRID:AB_400543 |
| Mouse monoclonal anti-CD11b, PC5, clone Bear1 | Beckman Coulter | IM3611; RRID:AB_131151 |
| Mouse monoclonal anti-CD13, APC, clone WM15 | BD | 557454; RRID:AB_398624 |
| Mouse monoclonal anti-CD14, BV605, clone M5E2 | BD | 564054; RRID:AB_2687593 |
| Mouse monoclonal anti-CD15, FITC, clone MMA | BD | 347423; RRID: AB_400299 |
| Mouse monoclonal anti-CD19, V450, clone HIB19 | BD | 560353; RRID:AB_1645564 |
| Mouse monoclonal anti-CD33, APC, clone P67.6 | BD | 340474; RRID:AB_400518 |
| Mouse monoclonal anti-CD33, BV786, clone WM53 | BD | 740974; RRID:AB_2740599 |
| Mouse monoclonal anti-CD34, APC-Cy7, clone 581 | BD | Custom conjugation |
| Mouse monoclonal anti-CD38, PE-Cy7, clone HB7 | BD | 335790; RRID:AB_399969 |
| Mouse monoclonal anti-CD38, BV421, clone HIT2 | BioLegend | 303526; RRID:AB_10983072 |
| Mouse monoclonal anti-CD41, PE-Cy5, clone P2 | Beckman Coulter | 6607116; RRID:AB_131057 |
| Mouse monoclonal anti-CD45, V500, clone HI30 | BD | 560777; RRID:AB_1937324 |
| Mouse monoclonal anti-CD45, AF700, clone HI30 | BD | 560566; RRID:AB_1645452 |
| Mouse monoclonal anti-CD45RA, FITC, clone HI100 | BD | 555488; RRID:AB_395879 |
| Rat monoclonal anti-CD49f, PE-Cy5, clone GoH3 | BD | 551129; RRID:AB_394062 |
| Mouse monoclonal anti-CD56, BV605, clone NCAM16.2 | BD | 562780; RRID:AB_2728700 |
| Mouse monoclonal anti-CD71, FITC, clone L01.1 | BD | 347513; RRID:AB_400316 |
| Mouse monoclonal anti-CD71, BV650, clone L01.1 | BD | 745273; RRID:AB_2742854 |
| Mouse monoclonal anti-CD90, APC, clone 5E10 | BD | 740585; RRID:AB_2740286 |
| Mouse monoclonal anti-CD90, APC, clone 5E10 | BD | 559869; RRID:AB_398677 |
| Mouse monoclonal anti-CD117, APC, clone 104D2 | BD | 333233; RRID:AB_2868677 |
| Mouse monoclonal anti-CD135 (FLT3), biotin, clone 4G8 | BD | 624008, Custom conjugation |
| Streptavidin PE | BD | 554061; AB_10053328 |
| Mouse monoclonal anti-GlyA, PE, clone KC16 | Beckman Coulter | IM2211U; RRID:AB_131213 |
| Mouse monoclonal anti-HLA-Dr, PC5, clone G46-6 | BD | 555813; RRID:AB_396147 |
| PE Mouse Anti-H2AX (pS139), clone N1-431 | BD | 562377; RRID: AB_2737611 |
| **Biological Samples** | | |
| Human fetal liver samples | Mount Sinai Hospital | N/A |
| Human umbilical cord blood samples | Trillium, William Osler and Credit Valley Hospitals | N/A |
| Human bone marrow samples | Centro Hospitalar Universitário de São João | N/A |
| Human acute myeloid leukemia samples | Princess Margaret Cancer Centre | N/A |
| **Chemicals, Peptides, and Recombinant Proteins** | | |
| G-CSF, human, premium grade | Miltenyi Biotec | 130-093-861 |
| TPO, human, premium grade | Miltenyi Biotec | 130-095-752 |
| IL-6, human, premium grade | Miltenyi Biotec | 130-093-932 |
| SCF, human, premium grade | Miltenyi Biotec | 130-096-695 |

| | | |
|---|---|---|
| IL-3, human, premium grade | Miltenyi Biotec | 130-095-069 |
| Propidium Iodide - 1.0 mg/mL Solution in Water | Thermo Fisher | P3566 |
| Propidium Iodide - 50 µg PI/ml Solution in PBS (pH 7.4) | BD | 556463 |
| AmpliTaq Gold 360 master mix | Thermo Fisher | 4398881 |
| Alt-R S.p. Cas9 Nuclease V3 | IDT | 1081059 |
| Collagenase IV | Stem Cell Technologies | 07909 |
| DNAse I | Roche | 11284932001 |
| DNase-Free RNase A, affinity purified, 1 mg/mL | Ambion | AM2271 |
| Proteinase K | Zymo Research | D3001220 |
| Ammonium Chloride | Stem Cell Technologies | 07850 |
| Bovine Serum Albumin Fraction V | Roche | 10735086001 |
| DMSO | Fisher Scientific | D128-500 |
| Agarose | Thermo Fisher | 16500500 |
| Formalin solution, 10% | Sigma | HT501128 |
| Sytox Blue | Thermo Fisher | S34857 |
| Giemsa stain, modified | Sigma | GS500-500ML |
| EcoRV (10U/uL) | NEB | R0195S |
| **Critical Commercial Assays** | | |
| CD34 MicroBead kit | Miltenyi Biotec | 130-046-702 |
| Mouse cell depletion kit | Miltenyi Biotec | 130-104-694 |
| Agencourt GenFind V2 | Beckman Coulter | A41497 |
| StemSep Human Progenitor Cells 10mL Kit | Stem Cell Technologies | 12470 |
| Anti-Human CD41 TAC | Stem Cell Technologies | 14060 |
| RNeasy Micro kit | Qiagen | 74004 |
| ZR-96 DNA clean-up kit | Zymo Research | D4018 |
| SMART-Seq V4 Ultra Low Input RNA kit | Clontech | 634891 |
| Nextera XT DNA Library Preparation kit | Illumina | FC-131-1096 |
| High Sensitivity DNA kit | Agilent | 5067-4626 |
| RNA 6000 Pico kit | Agilent | 5067-1513 |
| FITC Annexin V Apoptosis Detection Kit II | BD | 556570 |
| **Deposited Data** | | |
| Raw RNAseq data | This paper | EGA: under submission |
| Processed RNAseq data | This paper | GEO: GSE268962 |
| **Experimental Models: Organisms/Strains** | | |
| Mouse: *NOD.Cg-*$Prkdc^{scid}Il2rg^{tm1Wjl}$*/SzJ* (NSG) | The Jackson Laboratory | JAX: 005557 |
| Mouse: *NOD.Cg-*$Prkdc^{scid}Il2rg^{tm1Wjl}Kit^{em1Mvw}$*/SzJ* (NSG-W41) | Leonard D. Shultz | N/A |
| NOD.Cg-$Prkdc^{scid}$ $Il2rg^{tm1Wjl}$ Tg(CMV-IL3,CSF2,KITLG)1Eav/MloySzJ (NSG-SGM3) | The Jackson Laboratory | JAX:013062 |
| MS-5 (murine stroma cell line) | M.D. Minden's laboratory | N/A |
| **Oligonucleotides** | | |
| Alt-R CRISPR-Cas9 crRNA | IDT | N/A |
| Alt-R CRISPR-Cas9 tracrRNA | IDT | 1072534 |
| Alt-R Cas9 Electroporation Enhancer | IDT | 1075916 |
| Control (OR2W5) gRNA-1: GACAACCAGGAGGACGCACT | Wagenblast et al., 2019 | N/A |
| Control (OR2W5) gRNA-2: CTCCCGGTGTGGACGTCGCA | Wagenblast et al., 2019 | N/A |
| FLT3 gRNA-4 (exon 20): GATTCCAACTATGTTGTCAG | This paper | N/A |
| FLT3 gRNA-8 (exon 20): GGTGACAAGCACGTTCCTGG | This paper | N/A |

| | | |
|---|---|---|
| FLT3 gRNA-16 (exon 6): GCTTCATGAATTATTTGGGA | This paper | N/A |
| FLT3 gRNA-18 (exon 6): TGCCAGAAATGAACTGGGCA | This paper | N/A |
| KIT gRNA-4: GCTGAGCTTTTCTTACCAGG | This paper | N/A |
| Control (OR2W5) forward primer: 5’-TCGGCCTGGACTGGAGAAAA-3’ | Wagenblast et al., 2019 | N/A |
| Control (OR2W5) reverse primer: 5’-GAGACCACTGTGAGGTGAGA-3’ | Wagenblast et al., 2019 | N/A |
| ITD forward primer: 5’- GCAATTTAGGTATGAAAGCCAGC -3’ | Murphy *et al*, 2003 | N/A |
| ITD reverse primer: 5’- CTTTCAGCATTTTGACGGCAACC -3’ | Murphy *et al*, 2003 | N/A |
| TKD forward primer: 5’-GTAAAACGACGGCCAGCCGCCAGGAACGTGCTTG -3’ | Murphy *et al*, 2003 | N/A |
| TKD reverse primer: 5’-CAGGAAACAGCTATGACGATATCAGCCTCACATTGCCCC -3’ | Murphy *et al*, 2003 | N/A |
| FLT3_exon20 forward primer: 5′-CAGAAGCCGCACAAAGAACTG -3′ | This paper | N/A |
| FLT3_exon20 reverse primer: 5′-CTAGCTAGCCCAAGGACAGATG -3′ | This paper | N/A |
| FLT3_exon6 forward primer: 5′-CTCTGAGCTCTTAGCTGAAACCA -3′ | This paper | N/A |
| FLT3_exon6 reverse primer: 5′-TGCATTTACTGTGACCTGAAGGA -3′ | This paper | N/A |
| KIT forward primer: 5′- TAGGGTTGCAAGTGGGTGTTT -3′ | This paper | N/A |
| KIT reverse primer: 5′- GTCTATGCTTCACCCCAAGCC -3 | This paper | N/A |
| **Software and Algorithms** | | |
| FlowJo, 10.6.1 | BD | http://www.flowjo.com/ |
| GraphPad 9.0 | GraphPad Software | https://www.graphpad.com/ |
| CRoatan | Erard *et al*, 2017 | https://croatan.hannonlab.org/ |
| Benchling | Benchling | http://www.benchling.com/ |
| ICE Synthego | Conant *et al* 2022 | https://ice.synthego.com |
| DECODRv3.0 | Bloh *et al*, 2021 | https://decodr.org/ |
| ELDA | Hu *et al*., 2009 | http://bioinf.wehi.edu.au/software/elda/index.html |
| STAR 2.5.2b | Dobin et al., 2013 | N/A |
| EdgeR 4.0.16 | Robinson et al., 2010 | N/A |
| GSEA | N/A | https://www.gsea-msigdb.org/gsea/index.jsp |
| Cytoscape 3.10.2 | N/A | https://cytoscape.org/download.html |
| HTSeq 0.7.2. | Anders et al., 2015 | N/A |
| Leica Application Suite V3.3.0 | Leica | N/A |
| Aperio ImageScope | Leica | N/A |
| **Other** | | |
| IMDM media | Thermo Fisher | 12440053 |
| X-VIVO 10 hematopoietic serum-free culture media | Lonza | 04743Q |
| Parafdormaldehyde 32% aquos solution | Electron Microscopy Sciences | 15714-S |
| Perm Buffer III | BD | 558050 |
| BIT 9500 | Stem Cell Technologies | Cat #09500 |
| IMEM | Thermo Fisher | A1048901 |
| Fetal Bovine Serum (FBS) | GE Healthcare | SH3039603 |
| Fetal Bovine Serum (FBS) | Wisent | 185720 |
| Phosphate-Buffered Saline (PBS), pH 7.4 | Thermo Fisher | 10010049 |
| L-Glutamine | Thermo Fisher | 25030081 |
| Penicillin-Streptomycin | Thermo Fisher | 15140122 |

| | | |
|---|---|---|
| autoMACS Running Buffer | Miltenyi Biotec | 130-091-221 |
| Nuclease-free $H_2O$ | IDT | 11-05-01-14 |
| TE buffer | IDT | 11-01-02-02 |
| Lymphocyte Separation Media | Multicell | 305-010-CL |
| 4D-Nucleofector | Lonza | N/A |
| P3 primary cell 4D-nucleofector X Kit S | Lonza | V4XP3032 |
| P3 Primary Cell 4D-Nucleofector X Kit L | Lonza | V4XP-3024 |
| 96-well clear round bottom not treated plate | Corning | 351177 |
| Nunc 96-well flat bottom plates | Thermo Fisher | 167008 |
| 24-well TC plates | Falcon | 353047 |
| T75 suspension vented flasks | Sarstedt | 83.3911.502 |
| 96 well filter plate, 40µm | Pall | 8027 |
| 96 well PCR plate | Eppendorf | 951020362 |
| Magnetic stand | Thermo Fisher | AM10027 |
| Low-binding microcentrifuge tube, sterile | Axygen | MCT-175-C-S |
| LS column | Miltenyi Biotec | 130-042-401 |
| Shandon EZ Double cytofunnel | Thermo Fisher | A78710005 |
| 28gauge ½ cc syringe | Becton Dickinson | 329461 |
| 5ml syringe | BD | 309646 |
| 5ml tube with a 35µm cell strainer | Corning | 352235 |
| 40µm cell strainer | Corning | 431750 |
| 100µm cell strainer | Corning | 431752 |
| Razor blades | VWR | 55411-050 |
| FcR Blocking Reagent, human | Miltenyi Biotec | 130-059-901 |
| REact® 2 Buffer | Invitrogen | 16302-010 |
| Microscope Leica DM 4000B | Leica | https://www.leica-microsystems.com/products/light-microscopes/p/leica-dm4000-b/ |
